# Dendritic Morphology and Inhibitory Regulation Distinguish Dentate Semilunar Granule Cells from Granule Cells through Distinct Stages of Postnatal Development

**DOI:** 10.1101/2019.12.17.880005

**Authors:** Akshay Gupta, Archana Proddutur, Yun-Juan Chang, Vidhatri Raturi, Jenieve Guevarra, Yash Shah, Fatima S. Elgammal, Vijayalakshmi Santhakumar

## Abstract

Semilunar granule cells (SGCs) have been proposed as a morpho-functionally distinct class of hippocampal dentate projection neurons contributing to feedback inhibition and memory processing in juvenile rats. However, the structural and physiological features that can reliably classify granule cells (GCs) from SGCs through postnatal development remain unresolved. Focusing on postnatal days 11-13, 28-42, and >120, corresponding with human infancy, adolescence, and adulthood, we examined the somatodendritic morphology and inhibitory regulation in SGCs and GCs to determine the cell-type specific features. Unsupervised cluster analysis confirmed that morphological features reliably distinguish SGCs from GCs irrespective of animal age. SGCs maintain higher spontaneous inhibitory postsynaptic current (sIPSC) frequency than GCs from infancy through adulthood. Although sIPSC frequency in SGCs was particularly enhanced during adolescence, sIPSC amplitude and cumulative charge transfer declined from infancy to adulthood and were not different between GCs and SGCs. Extrasynaptic GABA current amplitude peaked in adolescence in both cell types and was significantly greater in SGCs than in GCs only during adolescence. Although GC input resistance was higher than in SGCs during infancy and adolescence, input resistance decreased with developmental age in GCs while it progressively increased in SGCs. Consequently, GCs input resistance was significantly lower than SGCs in adults. The data delineate the structural features that can reliably distinguish GCs from SGCs through development. The results reveal developmental differences in passive membrane properties and steady state inhibition between GCs and SGCs which could confound their use in classifying the cell types.

---

The dentate gyrus, the primary gateway for cortical inputs to the hippocampus, plays a unique role in memory processing as a center for sparse coding, mediated by the hyperpolarized resting membrane potential of granule cells as well as the robust synaptic and extrasynaptic GABAergic inhibition (Dengler and Coulter 2016). The classical dentate projection neurons, granule cells (GCs), which number in over a million in the rat brain, rest at more hyperpolarized potentials than most hippocampal neurons, receive powerful feedback inhibition, and project strong “detonator” synapses to CA3 enabling sparse yet reliable transmission (Dengler and Coulter 2016; Engel and Jonas 2005). Recent characterization of semilunar granule cells (SGCs), a dentate cell-type originally identified by Ramón y Cajal (1953), has revealed a second class of dentate projection neurons that are located in the inner molecular layer (IML), has distinctive somato-dendritic structure, persistent firing, and robust inhibition (Gupta et al. 2012; Larimer and Strowbridge 2010; Williams et al. 2007). While SGCs have been physiologically defined in rat (Gupta et al. 2012; Williams et al. 2007), cells with dendritic morphology consistent with SGCs have been observed in primates (Duffy and Rakic 1983; Seress and Frotscher 1990). Additionally, physiological recordings from cells with morphological features of SGCs have been performed in mouse and rabbit (Sancho-Bielsa et al. 2012; Save et al. 2018), indicating their presence across mammalian species. Due to their unique persistent firing in response to inputs, SGCs have been proposed to play a crucial role in regulating GC feedback inhibition, sparse coding of inputs, and pattern separation (Larimer and Strowbridge 2010; Walker et al. 2010). Recent studies suggest that SGCs are preferentially involved in memory engrams (Erwin et al. 2020). Since SGCs lack a cell-specific neurochemical marker, morphological characteristics, and lower input resistance (R_in_) are currently the primary approach to identify SGCs. However, the limited literature on SGC morphology and synaptic inputs during development have impeded further analysis of the role of SGCs in the dentate circuit.

Developmentally, both SGCs and GCs express the homeodomain transcription factor Prox1 (Gupta et al. 2012) and derive from the same neural precursor pool, although SGCs differentiate in a more restricted embryonic phase, demonstrating a common lineage between SGCs and GCs (Kerloch et al. 2018; Save et al. 2018). Based on the embryonic developmental analysis, SGCs are estimated at 3% of the neurons generated from the dentate neurogenic pool from which GCs derive (Save et al. 2018). Like GCs, SGCs have dendrites in the dentate molecular layer, and axons projecting to CA3. However, they can be distinguished from GCs based on their expansive dendritic span, large semi-lunar somata in the inner molecular layer (IML), and frequent presence of IML axon collaterals (Gupta et al. 2012; Williams et al. 2007). In earlier studies in rats, we defined SGCs on the basis of their wider dendritic angle (Gupta et al. 2012), which was subsequently confirmed in mice (Save et al. 2018). To date, morphometric analysis of SGCs has been limited to a narrow window of postnatal day (PD) 14-42 in rats, which is consistent with the neurological developmental state of human adolescence (Semple et al. 2013; Sengupta 2013), or in an embryonically labeled subgroup of neurons in young adult mice (Save et al. 2018). However, whether SGCs retain their distinct structural characteristics through postnatal development has not been examined. Recent studies have suggested layer specific differences in GC morphology (Kerloch et al. 2018) and disease-related changes in GC dendritic features (Freiman et al. 2011; Llorens-Martin et al. 2015; von Campe et al. 1997). However, without knowing the specific structural features that distinguish GCs from SGCs, it is difficult to interpret whether reports on changes in GC morphology reflect inclusion of structurally distinct SGCs in the datasets. Moreover, the ability to reliably distinguish SGCs from GCs across animal age is a prerequisite to elucidating the unique role that SGCs play in dentate processing. Unsupervised analysis of dendritic morphometric features is ideally suited to examine whether SGCs remain distinct from GCs through development and to elucidate the somato-dendritic features that specify the cell types.

Physiologically, SGCs show prolonged firing, reduced spike frequency adaptation, and lower R_in_ than GCs (Gupta et al. 2012; Save et al. 2018; Williams et al. 2007). Unlike GCs, SGCs respond to perforant-path stimulation with persistent firing lasting several seconds, which correlates with periods of increased hilar activity termed “up-states” (Larimer and Strowbridge 2010). SGCs have larger NMDA currents (Williams et al. 2007) than GCs, make synaptic contacts with hilar mossy cells and interneurons and have been proposed to drive granule cell feedback inhibition (Larimer and Strowbridge 2010). Apart from excitation, we previously demonstrated that SGCs have greater synaptic and extrasynaptic GABA currents (Gupta et al. 2012). Additionally, we identified that GCs and SGCs have diametrically opposite changes in synaptic and tonic GABA currents after brain injury suggesting that differences in inhibition which could contribute to distinct roles for SGCs and GCs in the dentate circuit. GCs undergo changes in both tonic and synaptic GABA currents during postnatal development (Hollrigel et al. 1998; Holter et al. 2010; Lee and Liou 2013). In parallel, studies on adult-born GCs labeled at specific time points show cell-age dependent maturation of GABA currents, ultimately reaching the levels similar to mature GCs in age-matched animals (Dieni et al. 2012; Li et al. 2012). Yet, whether inhibition in SGCs remains distinct from mature GCs through postnatal development is not known. While GCs show prolonged functional maturation (Yu et al. 2013a), age-dependent changes in inhibitory regulation of SGCs remains to be tested. This study was conducted to explicitly to determine the characteristic structural features which can be used to distinguish SGCs from GCs through postnatal developmental stages representing infancy (PD 11-13, prior to rodent eye opening), adolescence (PD 28-42 days, period of cortical maturation), and adult (> PD 120, cortical and sexual maturity) stages of development (Semple et al. 2013) and to identify the differences in steady-state inhibitory regulation of the two cell types through maturation of the dentate circuit.

## Materials and Methods

### Animals

All experiments were performed in accordance with IACUC protocols approved by Rutgers-NJMS, Newark, NJ, and the University of California at Riverside, CA and in keeping with the ARRIVE guidelines. The study included Wistar rats ranging in age from 11-13 days, 28-42 days, and 120-180 days designated as infancy, adolescence, and adulthood, respectively (Semple et al. 2013; Sengupta 2013). Due to the potential effects of hormonal variation on GABA currents, slice recordings in rats from >28 days were restricted to males. A subset of recordings was derived from surgical or saline-injected controls for an independent study and was pooled with data from naïve rats from which they showed no statistical difference.

### Slice Physiology

Rats were anesthetized with isoflurane and decapitated. Horizontal brain slices (300μm) were prepared in ice-cold sucrose artificial cerebrospinal fluid (sucrose-aCSF) containing the following (in mM): 85 NaCl, 75 sucrose, 24 NaHCO_3_, 25 glucose, 4 MgCl_2_, 2.5 KCl, 1.25 NaH_2_PO_4_, and 0.5 CaCl_2_ using a Leica VT1200S Vibratome. The low-sodium sucrose-aCSF has been shown to improve interneuron viability in adult slices (Tanaka et al. 2008). The slices were sagittally bisected and incubated at 32°C for 30 min in a submerged holding chamber containing an equal volume of sucrose-aCSF and recording aCSF, and subsequently were held at room temperature (22-23°C) prior to recordings. The recording aCSF contained the following (in mM): 126 NaCl, 2.5 KCl, 2 CaCl_2_, 2 MgCl_2_, 1.25 NaH_2_PO_4_, 26 NaHCO_3_, and 10 D-glucose. All solutions were saturated with 95% O_2_ and 5% CO_2_ and maintained at a pH of 7.4 for 1–6 h. Slices were transferred to a submerged recording chamber and perfused with oxygenated aCSF at 33°C. Whole-cell voltage-clamp recordings from putative GCs in the dentate granule cell layer (GCL) and SGCs in the inner molecular layer (IML) were performed using infrared differential interference contrast visualization techniques (Gupta et al. 2012; Yu et al. 2016) with a Nikon Eclipse FN-1 microscope, using a 40X, 0.80 NA water-immersion objective. Recordings were obtained at least 50μm below the slice surface in a field with several viable hilar neurons, typically within 1-6 hours after dissection in infant/adolescent mice and 1-3 hours following dissection in adult mice to ensure optimal slice health. Analysis was restricted to cells with resting membrane lower than −60 mV. Recordings were obtained using MultiClamp 700B (Molecular Devices). Data were low pass filtered at 3 kHz, digitized using Digidata 1440A, and acquired using pClamp10 at 10 kHz sampling frequency. Tonic and synaptic GABA currents were recorded in aCSF with no added GABA or GABA transporter antagonists in the recording solution (Gupta et al. 2012; Yu et al. 2013b). Voltage-clamp recordings of inward GABA currents were obtained from a holding potential of −70 mV using microelectrodes (5–7 MΩ) containing (in mM): 125 CsCl_2_, 5 NaCl, 10 HEPES, 2 MgCl_2_, 0.1 EGTA, 2 Na-ATP, and 0.5 Na-GTP, titrated to a pH 7.25 with CsOH. Biocytin (0.2%) was included in the internal solution for post hoc cell identification, and the glutamate receptor antagonist kynurenic acid (3 mM KynA, Tocris Bioscience) was included in the external solution to isolate GABA currents. Passive membrane properties including resting membrane potential and R_in_ were recorded in current clamp using an internal containing (in mM) 126 K-gluconate, 4 KCl, 10 HEPES, 4 Mg-ATP, 0.3 Na-GTP, 10 Phosphocreatine and 0.2% biocytin (Gupta et al. 2012). Among neurons recorded in the IML, only cells showing obtuse dendritic angles, presence of dendritic spines and axon projecting to the hilus (Gupta et al. 2012) (Supplementary Figure 1) were included in the physiological analysis as putative SGCs. The somata of all GCs included in the analysis were located in the GCL and spanned the entire extent of the GCL (Supplementary Figure 2). Neurons with small somata located in the subgranular zone, putative immature granule cells, were not included in the analysis. Recordings were discontinued if series resistance increased by >20%. Access resistance was not different between cell-types or between developmental groups and analysis of access resistance failed to reveal systematic differences recordings in between adolescent and adult slices (Access resistance in MΩ in GC and SGC combined, Adolescent 21.6±1.8 and Adult 20.9±1.7, p=0.8 by *t* test) indicating consistency in slice health. Following establishment of whole cell mode, baseline recordings were obtained for a minimum of five minutes in the presence of KynA prior to addition of GABA blockers. Recordings with excessive baseline fluctuations during the 3-5 minutes of recordings in KynA were discarded. All salts were purchased from Sigma-Aldrich (St. Louis, MO). Tonic GABA current, the steady-state current blocked by the GABA_A_R antagonist bicuculline methiodide (BMI, 100μM, Sigma-Aldrich) or gabazine (SR95531, 10 μM, Sigma-Aldrich), was measured as described previously (Gupta et al. 2012; Yu et al. 2013b) using custom macros in IgorPro7.0 software (WaveMetrics). Following physiological recordings, slices were fixed in 0.2mM phosphate buffer containing 4% paraformaldehyde at 4°C for 2 days. Biocytin staining was revealed using Alexa Fluor 594-conjugated streptavidin (Gupta et al. 2012; Swietek et al. 2016).

### Morphometry and hierarchical cluster analysis

Sections were visualized and imaged using a Zeiss LSM 510 confocal microscope with a 20X, 0.5 NA objective. Cells in which the dendritic trunks appeared severed at the slice surface were excluded from morphometric analysis. Cell reconstructions from confocal image stacks were performed using the directional kernels user-guided reconstruction algorithm in Neurolucida 360 (MBF Bioscience) followed by manual correction and validation in 3D. About 10-20 percent of each dendritic arbor was reconstructed manually. Neurolucida Explorer (MBF Biosciences) was used to extract non-nominal or non-ordinal somatodendritic morphological quantitative parameters (defined in Supplementary Tables 1 & 2) for use in statistical comparisons and hierarchical cluster analysis. A total of 42 projection neurons in which the dendritic arbors were fully reconstructed were analyzed. The software failed to quantify a subset of somatic parameters in 6 cells, and these cells were excluded from cluster analysis.

Data were tested for uniform distribution and each quantified variable was fit to the sum of two or more Gaussian functions and quality of fit determined using maximum likelihood analysis (MLA; v2test) to assess normal distribution of parameters within each cell type. Variables with a nonuniform distribution were used for subsequent cluster analysis. A total of 42 somato-dendritic parameters (Supplementary Tables 1 & 2) from 36 morphologically reconstructed neurons, including both features measured in Neurolucida Explorer from the 3D reconstructions and parameters measured manually in 2D rendering (Neurolucida 360, MicroBrightfield) were analyzed. Unsupervised clustering and principal component analyses of morphological properties were conducted within R version 3.5.0, using R package Cluster, FactoMineR, and factoextra by an investigator (Y-J C) blinded to cell types and age groups. Hierarchical clustering on the selected principal components (PCs) was performed using Ward’s Criterion with Euclidean distance to generate the dendrogram. The clustering partition was obtained from hierarchical clustering and improved with K-means method (Husson et al. 2010). Summary of morphological data and statistical analysis of morphometric parameters are presented in Supplementary Tables 3 and 4, respectively.

### Analysis and Statistics

Individual spontaneous inhibitory postsynaptic currents (sIPSCs) were detected using custom software in Igor-Pro7.0 (Gupta et al. 2012). The investigator (AG) was blinded to cell type during analysis. Events were visualized, and any “noise” that spuriously met trigger specifications was rejected. Cumulative probability plots of sIPSC amplitude and frequency were constructed using IgorPro by pooling an equal number of sIPSCs from each cell. Kinetics and charge transfer were calculated from the averaged trace of all accepted sIPSC events. Rise time was measured as the time for amplitude to change from 20 to 80 % of peak. Amplitude weighted τ_decay_ was calculated from a two-exponential fit to the IPSC decay. sIPSC charge transfer was calculated as the area under the curve of the baseline adjusted average sIPSC trace. The summed sIPSC charge transfer over one second was calculated as the product of the sIPSC charge transfer and sIPSC frequency for each cell. R_in_ was calculated as the slope of the current voltage plots with voltage response (averaged over the last 400 ms) to one second current injections from −200pA to −40 pA (in 40 pA steps) in each cell. Membrane time constant (τ_membrane_) was obtained from a single exponential fit to the voltage response to a −200pA current injection. Data are shown as mean±SEM (standard error of the mean) or median and interquartile range (IQR) where appropriate and presented in Supplementary Table 5. Kruskal-Wallis One Way Analysis of Variance on ranks was conducted on data which failed Shapiro Wilk test for normalcy or equal variance test. Two-way ANOVA (TW-ANOVA, SigmaPlot 12.3) and Bonferroni correction were used to test for statistical differences in tonic GABA currents. Summary data and details of the statistical tests used are included in Supplementary Tables 5-8. Sample sizes were not predetermined and conformed with those employed in the field. Significance was set to p<0.05, subject to appropriate Bonferroni correction. All custom macros for analysis and data sets will be made available on request upon publication of the study.

## Results

### Somato-dendritic characteristics distinguish SGCs from GCs throughout postnatal development

To examine whether SGCs are structurally distinct from GCs throughout postnatal development, we undertook electrophysiological recordings from putative GCs and SGCs in the GCL and IML respectively, and filled cells with biocytin to recover their morphology. As illustrated by morphological reconstructions obtained from confocal images of biocytin filled neurons (Fig. 1), putative GCs exhibit compact molecular layer dendritic arbors while putative SGCs recorded in the IML consistently exhibited wider dendritic spread. These differences in dendritic spread are maintained throughout the three developmental stages tested. Additionally, the X-Y plane projection of the reconstructed neurons revealed the characteristic crescent shaped somata of SGCs. Interestingly, the arborization in the z plane was not different between GCs and SGCs (in μm, GC: 73.6±8.7, n =15; SGC: 69.7±9.6, n=18, p>0.05 by Student’s *t* test). Additionally, the z plane span of SGCs was lower than their X-Y span indicating that SGCs have a flattened crescentic profile. less than the slice thickness. Figure 1 illustrates pseudo color rendering of dendritic arbors with segments assigned on the basis of branch order in 3D reconstructions in Neurolucida 360. Color coding of the dendritic segments revealed that, in addition to differences in dendritic span, SGCs differed from GCs in the extent to which individual dendritic trees branched and the number of segments in different dendritic orders. Moreover, reconstructions suggest that GCs have a dense distribution of dendritic arbors in a compact volume while SGCs appear to have sparsely distributed dendrites in a larger volume. Thus, while 3D reconstructions demonstrate that SGCs maintain a *qualitative* pattern of dendritic arborization distinct from GCs at all postnatal ages examined, they reveal additional morphometric features which differ between cell types indicating a need for comprehensive unsupervised *quantitative* analysis of dendritic structure.

**Figure 1.**
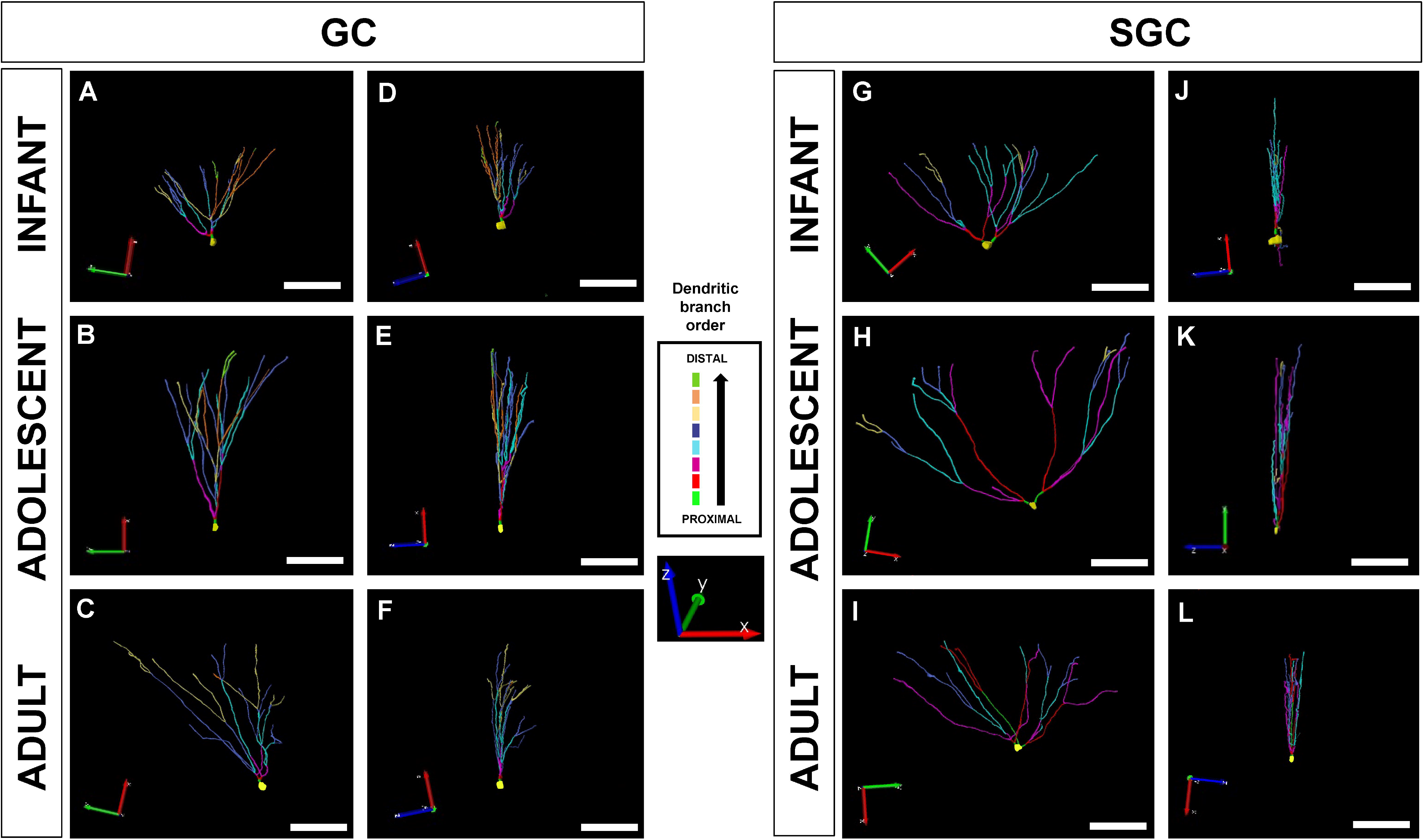
Somato-dendritic differences in dentate projection neurons - GCs and SGCs. Representative neuronal reconstructions of GC (1A-F) and SGC (1G-L) showing distinct dendritic arbors from infant (top), adolescent (middle) and adult (bottom) age groups. Note that images are shown in different planes for each age group. The complexity of dendrites is shown in XY planes for GCs and SGCs ‘A, B, C’ and ‘G, H, I’ respectively and perpendicular view is shown in ‘D, E, F’ for GCs and ‘J, K, L’ for SGCs. Inset images in the center represent the color coding for dendritic segments from proximal to distal axes. Scale bar: 100μm.

To determine if somato-dendritic parameters distinguish GCs from SGCs through development, we undertook the first unsupervised classification of dentate projection neurons on the basis of morphometry. Projection neurons were identified based on the presence of somata in the GCL (putative GCs) or IML (putative SGCs), axons with boutons entering the hilus and targeting CA3 (Supplementary Figure 1). As illustrated by the schematic of the somatic location of recorded cells (Supplementary Figure 2), GCs were recorded from the entire extent of the GCL. We find that both SGCs and GCs, have a high density of dendritic spines (Supplementary Figure 1), which we used as a criterion to distinguish projection neurons from local interneurons when the axon was not fully recovered. Morphometric parameters of the cells reconstructed in 3D were obtained from automated algorithms in Neurolucida 360 (definitions are included in Supplementary Tables 1 and 2). Principal Component Analysis (PCA) of 42 distinct morphometric parameters from 36 cells revealed a relatively high dimensional structure. The first three principal components (PCs) explained about 46% of the total variance in the data, while the first seven components retained over 85% of the variance (Supplementary Figure 3). PCA analysis of individual cells was projected on to the first three principal components and visualized in 3D representation with confidence interval (CI) ellipsoid set to 0.95, which suggested that the cells likely segregate by cell type (Fig. 2A) rather than developmental age of the rat (Fig. 2B).

**Figure 2.**
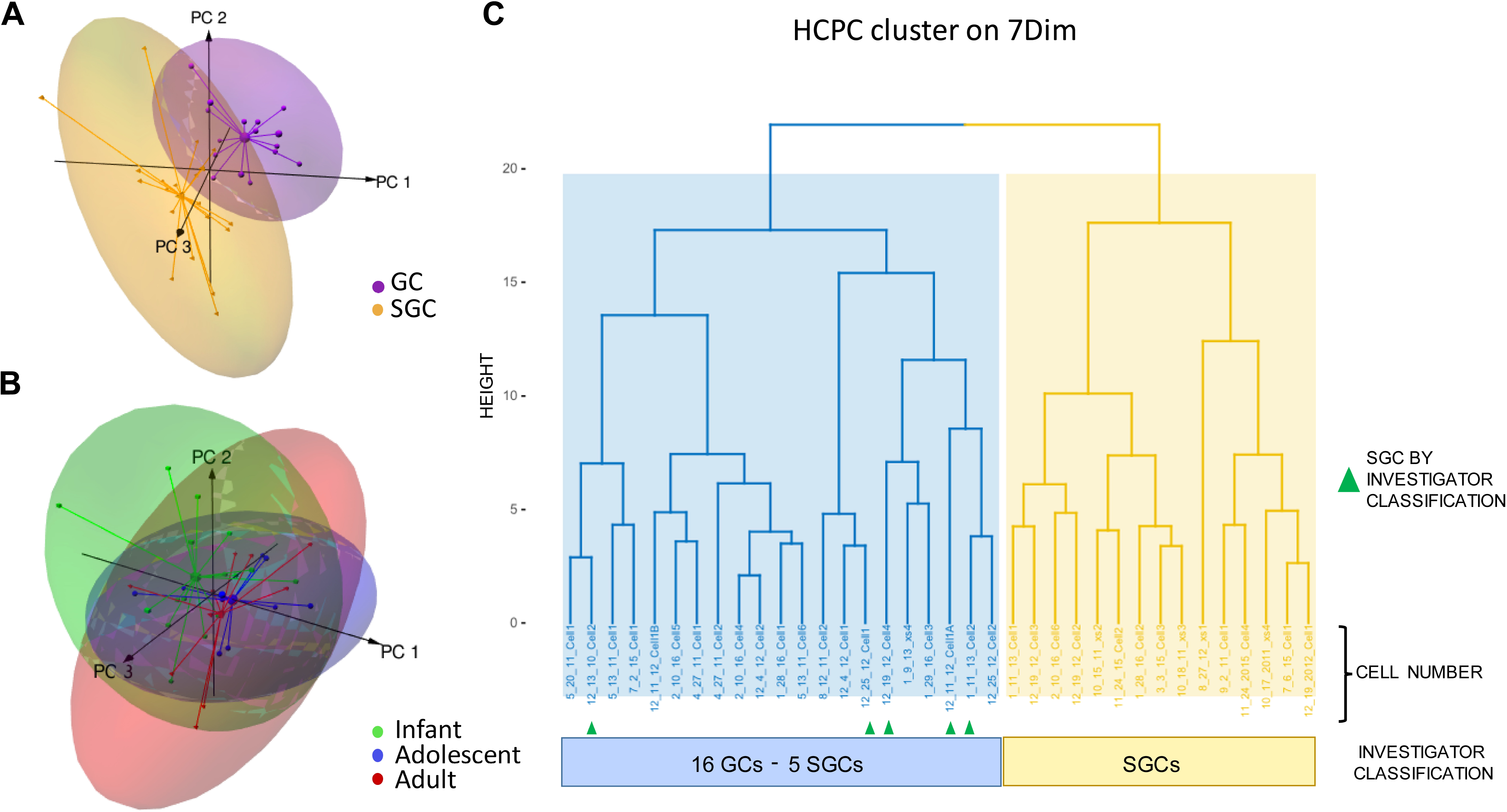
Hierarchical clustering on principal component reveals two major clusters of dentate projection neurons. (A-B) 3D representation of the principal component analysis of individuals resolved by first three principal components, with confidence interval (CI) ellipsoid set to 0.95. The plot suggests grouping by cell type (A) but not by age (B). (C) Hierarchical clustering on Principal Components based on 42 morphometric parameters (Supplementary Tables 1 & 2) was performed using Ward’s method with Euclidean distance to generate the dendrogram. Dendrogram classifies neurons into two putative clusters. Cells in which the classifier and experimenter failed to concur on classification are represented by green arrowheads based on investigator assigned classification.

Interestingly, although categorical variables including soma location were not included as a parameters for unsupervised analysis, clustering by PCA matched that of the investigator (Fig. 2). A total of 36 dentate projection neurons (16 cells in infant, 10 cells in adolescent and 10 cells in adult), recovered based on the quantitative morphological features, were included in the clustering analysis. A hybrid approach of hierarchical clustering on principal components (HCPC) which helps in de-noising multidimensional dataset was adopted (Husson et al. 2010). HCPC on the first seven components suggested two cluster partitioning groups (Fig 2C). Opening of the blinding revealed that the two putative clusters demonstrated a tendency for cells to cluster by putative cell-type. Cluster 1 included 16 cells classified as GCs and five classified as an SGC by the investigator. Clusters 2 consisted entirely of cell classified as SGCs by the investigator (A.G).

### Identity of principal morphometric parameters that best represent the PCs

The PCs were examined further to identify the top five morphological variables which best represent the PC (cos2 ≥ 0.7) and the morphological features that contributed to PC variability (Supplementary Figure 3). The number of first, second, and third order dendritic segments (note: number of first order dendritic segments equals number of primary dendrites), number of second order nodes and soma width where the top five morphological features that best represented the PCs while dendritic area and volume contributed most to variability (Supplementary Figure 3). Although the dendritic span and angle, used by the experimenter for classification (Gupta et al. 2012; Williams et al. 2007), contributed to the PC, morphology-based clustering revealed additional salient parameters including number of primary dendrites and dendritic segmentation which differed between clusters. Interestingly, one of the cells classified as SGCs by the investigator which clustered with putative GCs by unsupervised clustering was located in the IML (a feature that was not included in the cluster analysis), had a wide dendritic angle, but had two dendrites, with complex pattern of branching, typically seen in GCs, which could have driven the clustering with GCs (Supplementary Figure 2D_7_). Importantly, comparison of cell classification based on the unsupervised approach to that of the investigator (A.G.) confirmed that the investigator and PCA-based classifier agreed on the “grouping” of >85% (31 of 36) of cells examined. These findings confirm that GCs and SGCs are structurally distinct and demonstrate that the investigator can reliably discriminate the cell-types.

### Developmental changes in somato-dendritic morphology of dentate projection neurons

Using the investigator-assigned classifications, we next examined which specific morphological parameters showed cell-type and developmental differences. First, we focused on the parameters that contributed to PCs underlying cell classification. As predicted based on the PCA, the number of primary dendrites (first order segments), second order segments and nodes were different between SGCs and GCs, with SGCs having significantly more dendritic segments in the first three orders of dendrites (Fig. 3A, Supplementary Figure 4 & Supplementary Tables 3 & 4). However, the effect of *age* on the number of segments in each order was not statistically significant (Supplementary Figure 4A&B, Supplementary Tables 3 & 4). Similarly, soma width, soma aspect ratio and dendritic angle were greater in SGCs and failed to show age related changes (by TW-ANOVA; Fig. 3B&C, and Supplementary Tables 3 & 4). Thus, these parameters are ideally suited to distinguish cell-types regardless of age. In contrast, dendritic length showed a significant effect of age yet was not different between cell types (Fig. 3D, and Supplementary Tables 3 & 4). Of note, SGCs had significantly lower dendritic complexity than GCs and failed to show the age-dependent increase in complexity observed in GCs (Fig. 3E, and Supplementary Tables 3 & 4). Additional parameters that reflected 3D dendritic structure including convex hull surface area showed significant differences between cell-type and with age (Fig. 3F, Supplementary Tables 3 & 4). The convex hull surface area increased with development from infancy to adolescence and remained at adolescent levels in the adult (Fig. 3F, and Supplementary Tables 3 & 4) and appeared to contribute substantially to variability in the PCA analysis of pooled morphometric dataset (Supplementary Figure 3B). Finally, certain summed dendritic parameters including total numbers of dendritic terminals (ends), nodes, and segments showed neither cell-type nor age related differences (Supplementary Figure 4D-F, Supplementary Tables 3 & 4).

**Figure 3:**
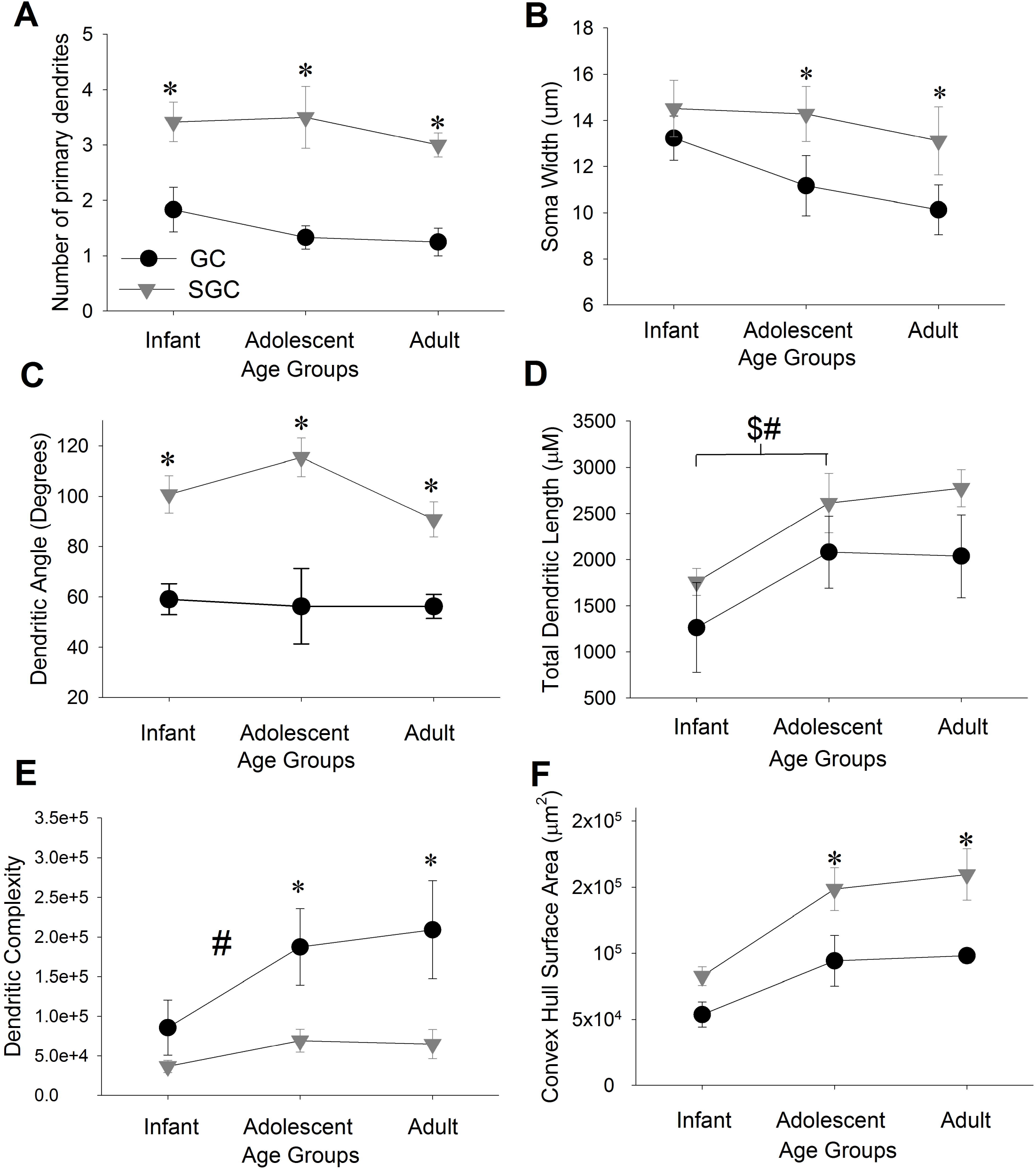
Comparison of morphometric parameters between GCs and SGCs at distinct developmental stages. Summary plots of averages no. of primary dendrites (A), soma width (B), dendritic angle (C), total dendritic length (D), dendritic complexity (E), and convex hull surface area (F) of GCs and SGCs at three developmental time points. *, #, and $ denote p<0.05 for differences between cell types, in GC across age groups and SGC across age groups, respectively by TW-ANOVA followed by post-hoc pairwise comparison (Supplementary Tables 3 & 4). N= GCs, 6 infant, 6 adolescent and 4 adult and SGCs, 9 infant, 5 adolescent and 6 adult.

### Developmental changes in spontaneous synaptic inhibition to GCs and SGCs

We previously demonstrated that SGCs from adolescent rats receive a greater frequency of action-potential driven sIPSCs than GCs from age-matched rats (Gupta et al. 2012). To determine if SGC inhibitory drive changes through postnatal development, we recorded sIPSCs from SGCs and GCs at three developmental stages. As illustrated in Fig. 4, SGCs consistently showed a higher frequency of sIPSCs compared to GCs from age matched rats. Both cell-types showed changes in sIPSC frequency with age. The frequency of sIPSCs in GCs increased from infancy through adolescence, peaked at adolescence and showed a slight, yet significant decrease in adults. (Fig. 4D, Supplementary Figure 5 and Supplementary Tables 5 & 6). Despite the decrease from adolescence to adulthood, the sIPSC frequency in GCs from adults was higher than that in infancy. SGCs, on the other hand, showed a distinct peak in sIPSC frequency during adolescence with a significant reduction in frequency in adults, back to the levels observed in infancy (Fig. 4D, Supplementary Figure 5, and Supplementary Tables 5 & 6). Thus, despite being consistently higher than in GCs, sIPSC frequency in SGCs appears to show a specific and transient enhancement during adolescence.

**Figure 4:**
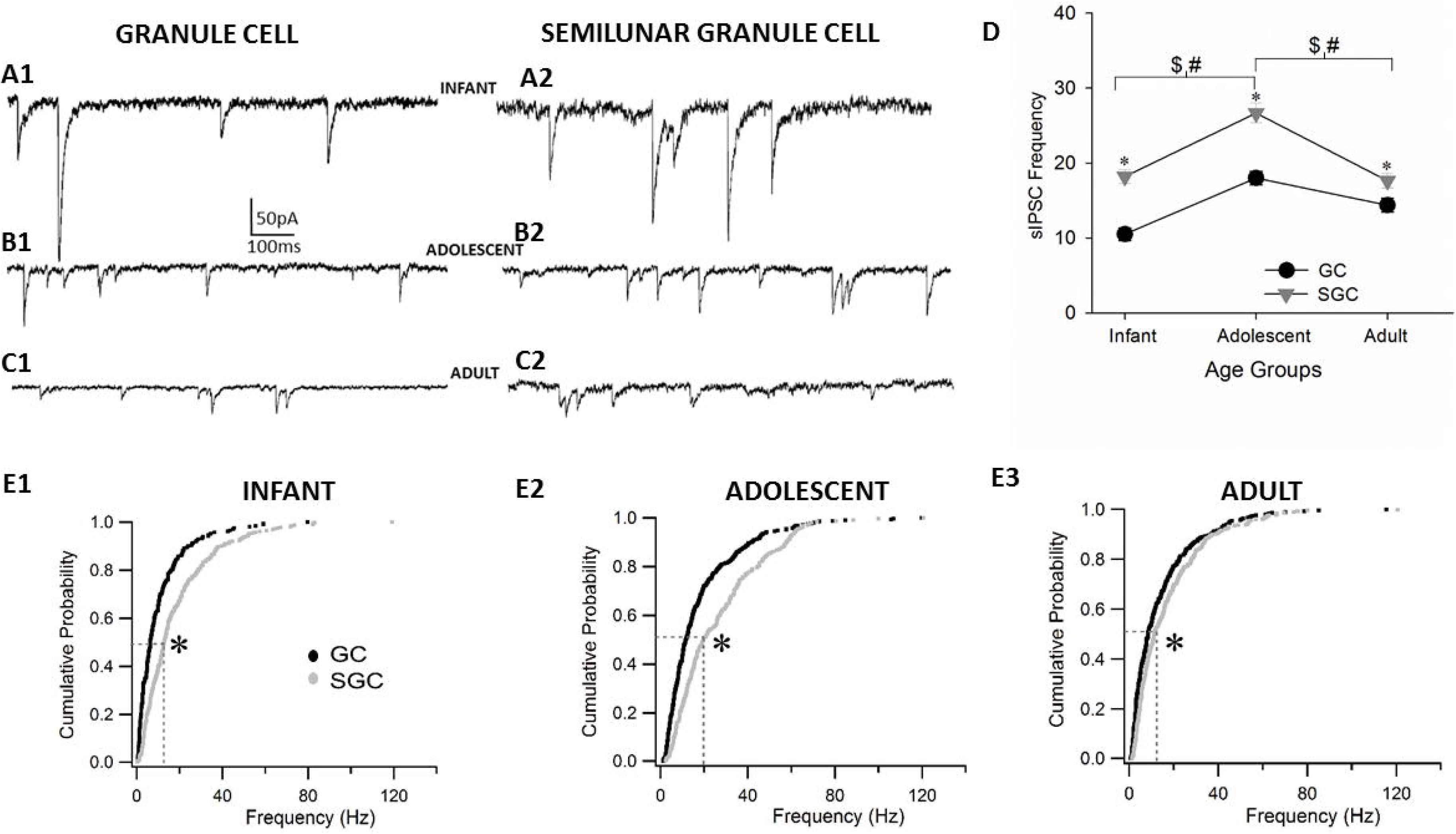
Developmental differences in sIPSC frequencies of GCs and SGCs. Representative sIPSC traces in GCs (left, A1, B1, C1) and SGCs (right, A2, B2, C2) in infant, adolescent, and adult age groups. Summary plot of developmental differences in sIPSC frequency between and within GCs and SGCs (D). *, #, and $ denote p<0.05 for differences between cell types, in GC across age groups and SGC across age groups, respectively by TW-ANOVA followed by post-hoc pairwise comparison (Supplementary Tables 5 & 6). Cumulative probability plots of sIPSCs frequency show differences between cell types at infant (E1), adolescent (E2) and adult (E3) age groups * denotes p<0.05 for differences between cell types by Kruskal-Wallis Test (Supplementary Tables 5 & 6).

Unlike sIPSC frequency, SGC sIPSC amplitude did not differ from age-matched GCs in infant and adolescent rats (Fig. 5C and Supplementary Tables 5 & 6), consistent with our earlier reports in adolescent rats (Gupta et al. 2012). However, in adult rats, sIPSC amplitude in GCs was larger than in SGCs (Fig. 5C). In GCs, sIPSC amplitude decreased significantly from infancy to adolescence and remained constant thereafter with differences in sIPSC between adolescence and adulthood not reaching statistical significance (Fig. 5C, Supplementary Tables 5 & 6). sIPSC amplitude in SGCs decreased through postnatal development with a significant reduction from infancy to adolescence, and a further decline from adolescence to adulthood (Fig. 5C, Supplementary Tables 5 & 6).

**Figure 5:**
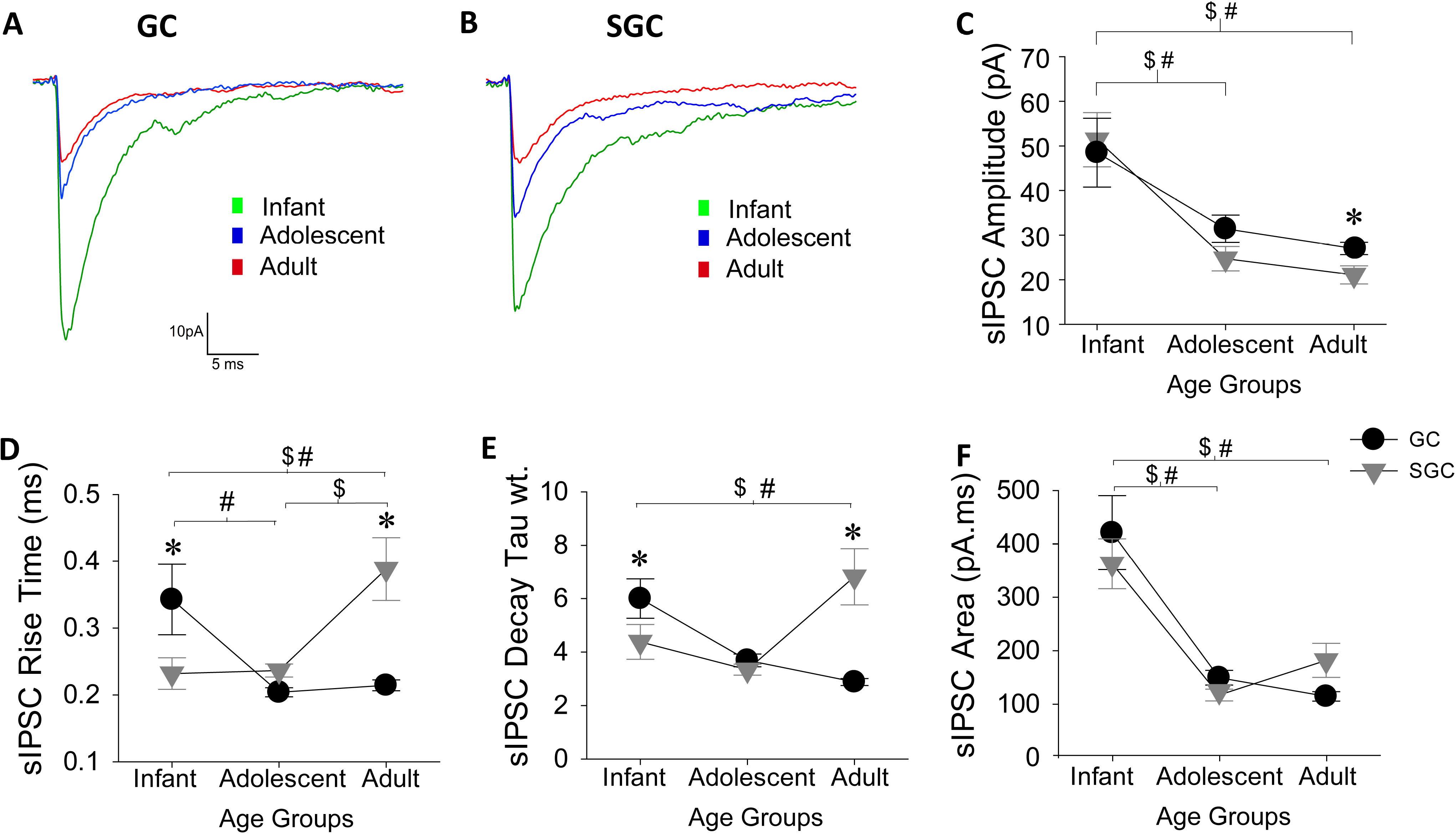
Developmental changes in average sIPSC parameters in GCs and SGCs. Overlay of representative average sIPSC waveforms from GCs (A) and SGCs (B) in different age groups. Summary plots of sIPSC amplitude (C), weighted τ_decay_ (D), 20-80 rise time (E), and charge transfer (F) at three developmental stages in both cell types. *, #, and $ denote p<0.05 for differences between cell types, in GC across age groups and SGC across age groups, respectively by TW-ANOVA followed by post-hoc pairwise comparison (Supplementary Tables 5 - 8).

Since distal inhibitory inputs can attenuate to a smaller amplitude at the soma (Soltesz et al. 1995), we sought to assess whether changes in the proportion of proximal versus distal dendritic synaptic inputs could contribute to developmental change in GC and SGC sIPSC amplitude. To determine if there is a systematic change in the amplitude of sIPSCs with age, we assigned sIPSCs to two groups based on their amplitudes: large and putative proximal and perisomatic (>50pA), and small (<50pA), potentially dendritic and calculated the proportion of these events classes in GCs and SGCs during development. Consistent with the developmental increase in dendritic length (Fig.3D), the proportion of large amplitude, presumed perisomatic events (>50pA) were highest in infants and reduced progressively with age while the smaller amplitude events <50pA events increased with age in both the cell types (Supplementary Figure 6). These findings suggest that developmental increase in dendritic length may contribute to decline in sIPSC amplitude over age in both cell types.

Systematic analysis of sIPSC kinetics identified an overall decline in τ_decay-WT_ in GCs with a significant reduction from infant to adolescent rats (Fig. 5D, Supplementary Table 5 & 7). Similarly, 20-80 rise time in GCs showed an overall decline with age with a significant reduction from infancy to adulthood (Fig. 5E, Supplementary Table 5 & 7). In contrast, both sIPSC 20-80 rise time and τ_decay-WT_ increased from infancy through adulthood in SGCs demonstrating a divergence in the effect of development on sIPSC kinetics between GCs and SGCs. Since the developmental changes in both sIPSC rise and τ_decay-WT_ showed parallel trends within a given cell type while differing between cell types, we evaluated whether developmental changes in cellular passive membrane parameters may underlie these changes. Consistent with the changes observed in sIPSC kinetics in GCs, both R_in_ and membrane time constant (τ_membrane_) trended to decline from infancy through adulthood with the decrease in R_in_ reaching statistical significance (Supplementary Fig. 7, Supplementary Tables 5 & 8). As with sIPSC kinetics in SGCs, both R_in_ and τ_membrane_ increased significantly from infancy through adulthood (Supplementary Fig. 7, Supplementary Tables 8). While direct correlation of the sIPSC kinetics with τ_membrane_ was not feasible due to use of different cohorts of recordings for sIPSC and passive parameter data, the results suggest that developmental changes in membrane passive properties could, among other factors, contribute to shift in sIPSC kinetics in cell types with postnatal development. Moreover, the data demonstrate that postnatal development has an opposite effect on R_in_ in GCs and SGCs. R_in_ in SGCs was lower than in GCs in infant and adolescent rats, a feature that has been used to distinguish the cell types (Erwin et al. 2020). However, in adult rats, R_in_ in SGCs was significantly higher than age matched GCs (Supplementary Fig. 7, Supplementary Tables 5 & 8) revealing that a lower R_in_ cannot consistently distinguish SGCs from GCs across all developmental time points.

Given the developmental and cell-specific changes in sIPSC peak amplitude and kinetics, we examined whether GCs and SGCs showed changes in sIPSC charge transfer during postnatal development. Consistent with changes in peak amplitude, sIPSC charge transfer declined with development in both cell types (Fig. 5F, Supplementary Tables 5 & 7). Despite the changes in kinetics, the net charge transfer was not different between GCs and SGCs from age-matched rats (Fig. 5F, Supplementary Tables 5 & 6). To comprehensively assess spontaneous synaptic inhibition in GCs and SGCs during development in a manner that would include divergent changes in frequency amplitude and kinetics between cell-types and during development, we estimated the cumulative sIPSC charge transfer over one second as a product of the sIPSC frequency and charge transfer for each cell. Interestingly, the cumulative sIPSC charge transfer over one second was maximum during infancy and declined with development in both GCs and SGCs (Supplementary Fig. 5C, Supplementary Table 5 & 7). Moreover, the cumulative sIPSC charge transfer over one second was not different between cell-types at any developmental stage examined (Supplementary Fig. 5C, Supplementary Table 5 & 7).

### Extrasynaptic GABA currents in SGCs peak during adolescence

Apart from GABAergic synaptic inputs, dentate GCs are known to express extra- and peri-synaptic GABA_A_ receptors that contribute to steady-state tonic GABA currents (Stell et al. 2003). We previously demonstrated the presence of tonic GABA currents in SGCs and identified that the amplitude of tonic GABA current in SGCs was greater than in age-matched adolescent GCs (Gupta et al. 2012). Here we find that although tonic GABA current amplitude in SGCs, measured as the baseline currents blocked by a saturating concentration of GABA_A_ receptor antagonists, was significantly greater than in GCs during adolescence, tonic GABA_A_ current amplitude was not different between GCs and SGCs during infancy or adulthood (Fig. 6). Both GCs and SGCs showed a significant increase in tonic GABA currents from infancy to adolescence which returned back to pre-adolescent levels in adults (Fig. 6 and Supplementary Tables 5 & 6).

**Figure 6:**
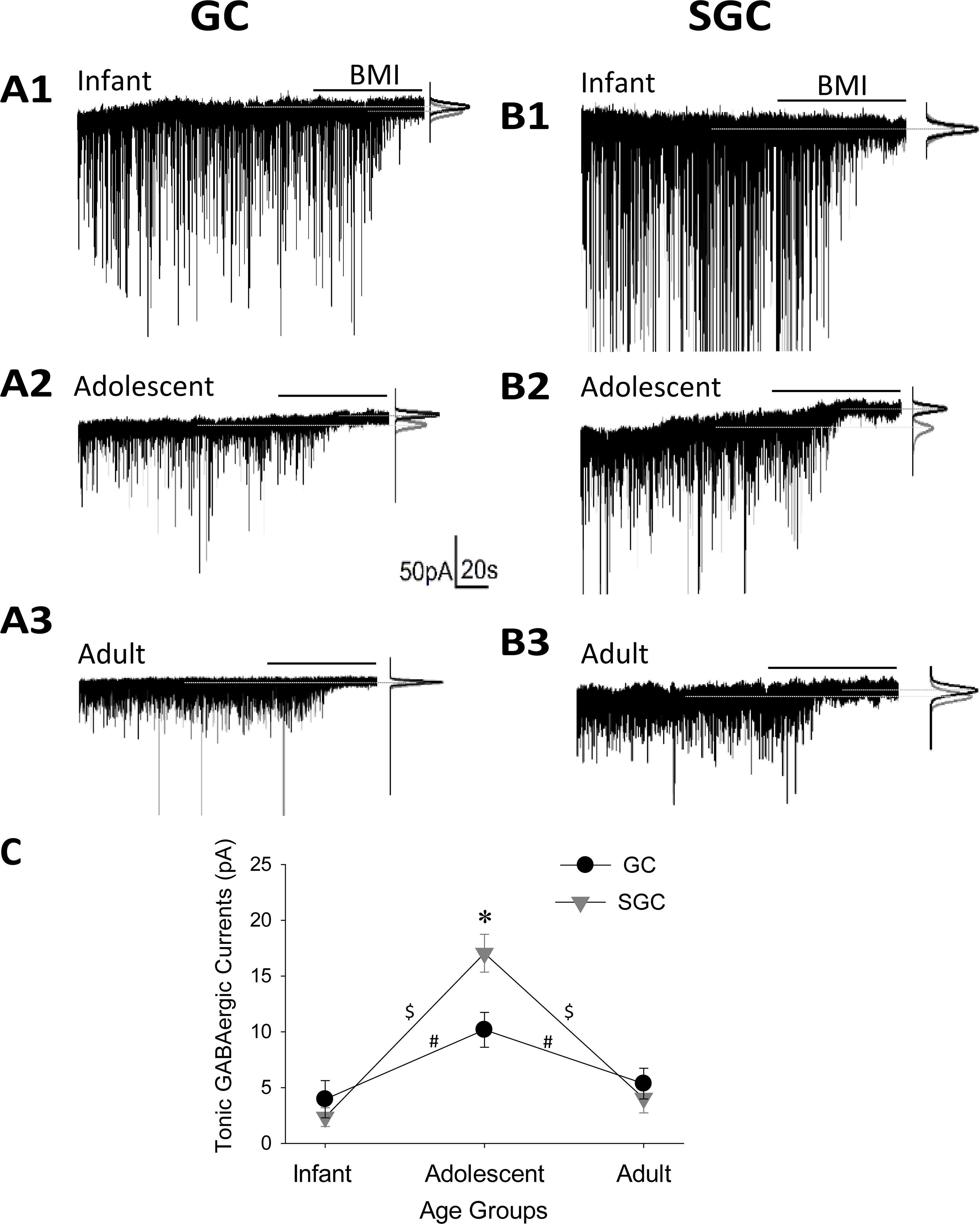
Extrasynaptic tonic GABAergic currents in SGCs peak during adolescence. Representative baseline current recordings in GCs (left, A1-3) and SGCs (right, B1-3) in infant, adolescent and adult age groups. Tonic GABA current is measured as the difference in baseline current upon perfusion of the GABA receptor antagonist BMI (100μM). Gaussian fit to the positive half of the baseline current, under basal conditions and in BMI, is illustrated on the right of each trace and was used to quantify tonic GABA. (C) Summary plot showing average tonic GABAergic currents at three distinct time points. *, #, and $ denote p<0.05 for differences between cell types, in GCs across age groups and in SGCs across age groups, respectively by TW-ANOVA followed by post-hoc pairwise comparison (Supplementary Tables 5 & 6).

## Discussion

Contemporary literature on the dentate gyrus describes the projection neurons as a largely homogeneous population of GCs with limited diversity (Kesner 2018). Since GCs are a unique subset of neurons that undergo neurogenesis and maturation through adulthood, structural and functional diversity in GCs is largely attributed to the maturation state of neurons within the circuit (Toda and Gage 2018). At any given postnatal developmental stage of the animal, immature GCs tend to be located closer to the hilar border of the cell layer, have less elaborate dendritic structures and are functionally more excitable than their mature counterparts (Kerloch et al. 2018; Overstreet-Wadiche and Westbrook 2006). However, emerging recognition of a structurally and functionally distinct subset of dentate projection neurons challenges the prevailing view that dentate projection neurons are a homogenous class of cells distinguished by developmental stages (Gupta et al. 2012; Save et al. 2018; Williams et al. 2007). SGCs, which have been characterized in the IML of the dentate gyrus, differ from GCs in their expansive dendritic arbors and show more sustained firing activity (Larimer and Strowbridge 2010; Williams et al. 2007). These and other features, including their distinct window of embryonic day 12-15 for SGC development (Save et al. 2018), enhanced excitatory drive (Larimer and Strowbridge 2010), and distinct synaptic and extrasynaptic inhibitory currents (Gupta et al. 2012) indicate that SGCs are distinct from GCs. Although SGCs have been proposed to sculpt feedback inhibition of GCs, gate dentate activity, and contribute to memory processing (Larimer and Strowbridge 2010; Walker et al. 2010), there is limited information on the structural and physiological differences between GCs and SGCs across postnatal development of the animal. This information is needed to determine which features consistently and reliably distinguish the cell types at all age groups. Our detailed and objective morphometric analysis conducted at three distinct developmental stages, namely, infant, adolescent, and adult age groups demonstrates that clustering dentate projection neurons based on somato-dendritic morphology distinguishes them into different “subtypes” corresponding to SGCs and GCs. Additionally, SGCs consistently maintained higher frequency of inhibitory synaptic inputs than GCs at all ages. Our results demonstrate that although both GCs and SGCs exhibit peaks in frequency of synaptic inhibitory inputs and magnitude of tonic GABA currents during adolescence, these parameters were significantly higher in SGCs than in GCs at this time point. However, the peak sIPSC amplitude and cumulative charge transfer were highest in infants, decreased with development and were not different between cell types. These findings demonstrate that SGCs are a structurally and functionally distinct subtype of dentate projection neurons which is in keeping with the emerging recognition of subpopulations among hippocampal and entorhinal projection neurons (Pilli et al. 2012; Soltesz and Losonczy 2018). Moreover, the significantly heightened tonic inhibition in SGCs during adolescence suggests that SGC activity could be more strongly modulated than GCs by a variety of neuroactive compounds including alcohol and neurosteroids, which selectively augment extrasynaptic GABA currents (Maguire and Mody 2009; Mody et al. 2007). Future studies examining effects of alcohol and neurosteroids on SGC function during adolescence coupled with analysis of developmental changes in excitatory drive and active properties of GCs and SGCs can provide a more comprehensive understanding of the network role of SGCs.

In addition to using unbiased approaches to classify SGCs as a distinct neuronal class, our data identify key age-invariant features to distinguish the SGCs from GCs. We find that the number of primary dendrites, dendritic angle, soma width and soma ratio are significantly higher in SGCs and can be used to categorize SGCs and GCs. In particular, the multiple primary dendrites observed in SGCs stands in stark contrast to the typical one to two apical dendrites observed in granule cell reconstructions (Thind et al. 2008). Indeed, the striking >85% correspondence between the unsupervised clustering and investigator assigned clustering likely stems from the investigator’s use of dendritic angle, soma width, which are significant contributors to the first PC, in addition to soma location to classify cell types. Dendritic length, on the other hand, increased during development but was not different between cell types indicating that processes reflecting developmental maturation are common to the cell types. Consistent with the presence of larger dendritic angle, the convex hull 3D volume and 3D surface area of SGCs were greater than that of GCs. However, there was also a developmental increase in these parameters from infancy to adolescence and a further stabilization into adulthood in both cell types, likely reflecting the developmental increase in hippocampal volume. While extensive precautions were taken to exclude cells with severed dendrites from morphometric analysis, it remains possible that the fills in slices may have missed some dendritic branches. However, our finding that the z axis thickness is not different between GCs and SGCs and is considerably lower than both slice thickness and X-Y span indicate that dendritic arbors of both cell types are likely fully represented within the slice. Interestingly, although the number of terminal nodes was not different, the dendritic complexity was significantly lower in SGCs than in GCs demonstrating differences in branching patterns. Difference in branching patterns including dendritic complexity can impact neuronal firing and intracellular signaling, as has been demonstrated in modeling studies (Li et al. 2015; van der Velden et al. 2012; van Elburg and van Ooyen 2010). Whether dendritic morphology can account for differences in intrinsic physiology between the dentate projection neuron types (Gupta et al. 2012; Save et al. 2018; Williams et al. 2007) remains to be examined. The other aspect where distinguishing cell type based on morphology becomes critical is in disease. Dentate granule cells are known to undergo changes in dendritic structure including alterations in complexity under physiological conditions and in trauma and neurodegenerative diseases (Llorens-Martin et al. 2015; Redila and Christie 2006; Villasana et al. 2015). The ability to distinguish GCs and SGCs across multiple developmental stages would be crucial to quantifying and interpreting changes in morphology in trauma, epilepsy, and neurodegenerative disease.

While there are clear differences in the structure, SGCs and GCs share several characteristics. Both SGCs and GCs are projection neurons, with dendrites in the dentate molecular layer and axons projecting to hippocampal CA3 (Gupta et al. 2012; Save et al. 2018; Williams et al. 2007). Several dendritic parameters including total numbers of dendritic segments, nodes, terminals, and dendritic tortuosity (Supplementary Table 3) showed neither cell type nor age-related differences. Similarly, SGCs, like GCs, have dendritic spines which can aid in distinguishing them from inhibitory neurons. Additionally, hilar axon collaterals of SGCs have “mossy fiber boutons” typically attributed to GCs (Supplementary Fig. 1 and Save et al. 2018). Notably, in pilot clustering analysis which included a few molecular layer inhibitory neurons, the interneurons clustered on a different branch of the dendrogram than SGCs and GCs (data not shown). The similarities between GCs and SGCs are not surprising, since we previously identified that SGCs express the Prospero homeobox protein 1 (Prox1) present in GCs indicating a shared lineage (Gupta et al. 2012). This shared lineage was confirmed by a recent study which identified a narrow developmental window of embryonic day 12-15 during which SGCs are produced from the same precursor niche as GCs (Save et al. 2018). Yet, the morphology of embryonically labeled SGCs were distinct from that of GCs labeled on the same day (Save et al. 2018), demonstrating that they are a distinct population of cells rather than a cohort of GCs with a different maturation state. In this context, studies examining dendritic properties of GCs with somata located in the outer third of the molecular layer have consistently reported wider dendritic fields and distinct dendritic arbors consistent with the possibility that the outer third of the granule cell layer may consist of a mixed population of GCs and SGCs (Green and Juraska 1985; Kerloch et al. 2018; Sun et al. 2013). Whether specific genetic, molecular or developmental cues guide the development of SGCs during neurogenesis or whether local molecular and spatial factors in the dentate GCL-molecular layer border contribute to elaboration of distinct dendritic arbors remains to be determined (Hatami et al. 2018; Lefebvre et al. 2015). Similarly, whether SGCs and GCs have comparable dendritic length distribution within the distinct axonal projection zones, namely, commissural/associational pathway in the IML, medial perforant path in the middle molecular layer and lateral perforant path in the outer molecular layer was outside the scope of the current study and merits further analysis. Although the mechanisms underlying specification of SGCs as a distinct population and the layer specific inputs to SGCs need further investigation, our objective unsupervised morphometric analysis identifies the key somato-dendritic structural features that distinguish SGCs from GCs through postnatal development.

A defining feature of the dentate gyrus is the presence of heavy inhibitory regulation (Coulter and Carlson 2007). Dentate GCs receive synaptic inhibition from a diverse population of neurons and are also under steady-state, tonic extrasynaptic inhibition (Coulter and Carlson 2007; Ewell and Jones 2010; Harney and Jones 2002; Stell et al. 2003). In an earlier study, we identified that SGCs are under stronger inhibitory regulation than GCs with higher frequency of inhibitory synaptic currents and greater amplitude of tonic GABA currents (Gupta et al. 2012). Here, we find that SGCs continue to receive greater spontaneous synaptic inhibitory events than GCs through postnatal development. This contrasts with the lack of difference in the frequency of spontaneous IPSCs in mature granule cells regardless of whether they were born embryonically or in adults (Laplagne et al. 2007) in spite of the complex and extended maturation of inhibitory synapses on to adult born granule cells (Dieni et al. 2013; Groisman et al. 2020). Interestingly, while the frequency of sIPSCs peaked during adolescence in both cell types, the amplitude decreases progressively with development. Additionally, while synaptic membrane kinetics decreased with development in GCs, they increased with development of SGCs. The combined effect of cell specific and developmental changes resulted in an overall decrease in cumulative synaptic inhibitory charge transfer from infancy to adulthood while maintaining similar charge transfer between age-matched GCs and SGCs. In addition to developmental changes in τ_membrane_ which could contribute to cell-specific regulation of synaptic kinetics, the roles of changes in GABA receptor subunits, dendritic pruning and synaptic distribution need to be considered in future works. Although lower R_in_ in SGCs has been considered a defining distinction from GCs (Erwin et al. 2020; Gupta et al. 2012; Williams et al. 2007), our demonstration that SGCs in adult rats have higher R_in_ than in GCs suggests that R_in_ may not be an age-invariant feature for cell classification.

Can dendritic structural features explain the cell-type specific and developmental differences in sIPSC frequency in dentate projection neurons? The consistently higher sIPSC frequency in SGCs is surprising as the dendritic lengths and location are not different between GCs and SGCs (Fig. 3). The wider dendritic distribution of SGCs, also reflected in the greater convex hull 3D volume, could allow for inputs from a larger group of inhibitory neurons to impinge on SGC dendrites, while GCs with their compact dendritic distribution may receive fewer inputs. It is also possible that the relatively early embryonic development of SGCs increases the chance of SGCs to receive more synaptic inputs compared to GCs which develop later into adulthood. However, since both dendritic length and convex hull 3D volume increase with postnatal development of both cells, changes in dendritic morphology and embryonic development are unlikely to account for the peak in sIPSC frequency in adolescence followed by decline in adults. The possibility of developmental increase in synapses from infancy to adolescence followed by pruning or synapse elimination into adulthood (Riccomagno and Kolodkin 2015; Tran et al. 2009) may be considered in future works. Although multiple precautions were taken to maintain tissue health in slices from adult rats, it is possible that differences in interneuron viability in slices from adolescent versus adult rats also contributed to decline in IPSC frequency with age. The progressive decrease in sIPSC amplitude with development in both SGCs and GCs could reflect the increase in dendritic length with development. Consistent with this proposal, the proportion of large amplitude events decrease progressively with development in both cell types (Supplementary Figure 6).

In parallel, the amplitude of extrasynaptic GABA currents in SGCs peaked during adolescence. Similarly, tonic GABA currents in GCs showed a peak during adolescence confirming the developmental increase reported in previous studies (Holter et al. 2010). Increase in expression of the extrasynaptic GABA_A_R δ subunit in both GCs and SGCs (Gupta et al. 2012; Maguire and Mody 2009) is likely to mediate the increase in tonic GABA currents from infancy to adolescence. Additionally, changes in synaptically released GABA accompanying changes in sIPSC frequency, reported here, could contribute to the peak in extrasynaptic GABA exhibited in adolescence. An interesting feature of tonic GABA currents mediated by GABA_A_R δ subunit is their robust enhancement by neurosteroids, raising the possibility that increases in ambient neurosteroids during adolescence (Harden and MacLusky 2004; Maguire and Mody 2009) could augment tonic GABA currents. Since tonic GABA currents mediated by GABA_A_R δ subunits are exquisitely sensitive to ethanol (Mody et al. 2007), the greater magnitude of tonic GABA currents in SGCs during adolescence is likely to render SGCs vulnerable to ethanol modulation and impact their role in dentate processing. Indeed, behavioral deficits following alcohol administration have been shown to be particularly accentuated during adolescence (Lacaille et al. 2015; Spanos et al. 2012). It would be important for future studies to ascertain how the differences in basal synaptic inhibition impact the recruitment and circuit function of SGCs and GCs during network activity and with development. Since basal inhibition in the dentate gyrus regulates synaptic plasticity and pattern separation at the level of the circuit (Dengler and Coulter 2016; Madar et al. 2019), and SGCs have been proposed to contribute to dentate pattern separation (Larimer and Strowbridge 2010), developmental changes in GC and SGC inhibition are likely to influence memory and cognitive performance.

Together, the structural and functional data identify SGCs as a cell type which differs from GCs in somato-dendritic structure and developmental plasticity of inhibitory inputs, extrasynaptic inhibition and membrane kinetics. Our data demonstrate that SGCs have heightened extrasynaptic inhibition during adolescence which would make them susceptible to endogenous and exogenous modulation of activity levels during adolescence. Notably, our results delineate salient structural features that enable anatomical identification of this subpopulation of dentate projection neurons. The novel data defining the structural features of SGCs will allow for future targeted analysis of their molecular profile and microcircuit connectivity to better understand their role in circuit function and behaviors. In conclusion, the fundamental characterization of SGCs presented here will support incorporation of SGCs into current models of the dentate gyrus and consideration of their role in dentate microcircuit processing in health and disease.

## Acknowledgements

We thank Dr. Luke Fritzky at the Rutgers Imaging Core for help with imaging and Dipika Sekhar for data entry. We thank Drs. Deepak Subramanian and Kelly A. Hamilton for thoughtful discussions and comments.

## Supplementary Figure legends

**Supplementary Figure 1: Representative images of a GC and SGC.** Images of a typical GC (A) and SGC (B) illustrate the somatic location, dendritic arbor, with high density of spines (insets, white arrows) and axons with boutons (white arrow heads) targeting CA3 used by experimenter to classify SGCs and GCs. Scale bar: 100μm; Inset scale bar: 20 μm.

**Supplementary Figure 2:** Distribution of somatic location of cells included in morphometric analysis. A. Schematic of the somatic location in the GCL (green box) and IML (orange box) overlaid on the cluster analysis distribution illustrates SGCs in blue and GCs in green. B_1-3_: Montage of biocytin filled images of representative GCs (labeled as B_1-3_ in A) with GCL demarcated by dotted lined identified based on Prox-1 (B_1-2_) or parvalbumin (B_3_) immunostaining. C. Representative SGC reconstruction in IML overlaid on image of the slice. D_1-7_. Examples of biocytin filled and reconstructed GCs and SGCs labeled as D_1-7_ in panel A. Note that the SGCs illustrated here clustered with putative GCs and are included in the white dotted area in A. All images were obtained a 20X magnification and reconstructed. Note the difference in soma shape of cells classified as GCs (D_1-3_) versus those classified as SGCs(D_4-7_) by investigator. D_7_. Note the increased dendritic complexity compared to putative SGCs in D_4-6_. GCL: Granule Cell Layer, IML: Inner Molecular Layer.

**Supplementary Figure 3: Analysis of Morphological Variables underlying Principal Components.** A. Histogram illustrates percentage of information retained by each dimension (principal component, PC). B. Factor maps illustrate the quality of representation of the morphometric variables measured by cos2 (square cosine, squared coordinates). The darker color indicates stronger contribution to variability to PC. C. Representation of the top five variables in the first two dimensions..

**Supplementary Figure 4: Comparison of morphometric parameters between GCs and SGCs at distinct developmental stages.** Summary plot of number of second (A) and third (B) order dendritic segments, second order nodes (C), total number of dendritic ends (D), total dendritic nodes (E) and total dendritic segments (F) in GCs and SGCs at the three age groups examined. * denotes p<0.05 for differences between cell types by TW-ANOVA followed by post-hoc pairwise comparison (Supplementary Tables 3 & 4). N= GCs, 6 infant, 6 adolescent and 4 adult and SGCs, 9 infant, 5 adolescent and 6 adult.

**Supplementary Figure 5: Cumulative plots of sIPSC frequency in GCs and SGCs through development.** Cumulative plots of sIPSC frequency comparing three age groups * denotes p<0.05 for differences between cell types by Kruskal-Wallis Test (Supplementary Table 6) in GCs (A) and SGCs (B). Summary plot of cumulative charge transfer over one second (C) at three developmental stages in both cell types. #, and $ denote p<0.05 for differences in GC across age groups and SGC across age groups, respectively by TW-ANOVA followed by post-hoc pairwise comparison.

**Supplementary Figure 6: Developmental differences in distribution of sIPSC amplitudes in GCs and SGCs.** Pie chart distributions illustrating subjective percentage distribution of high peak amplitude (>50pA) and smaller peak amplitude (<50 pA) amplitude sIPSCs in GCs (top) and SGCs (bottom) across age groups.

**Supplementary Figure 7: Analysis of developmental changes in passive membrane properties of GCs and SGCs.** Representative voltage traces in response to a −200pA current injection for one second in GCs and SGCs at the developmental stages under investigation reveal differences in R_in_ (A). Summary of R_in_ in the cell types (B). Summary plot of membrane time constant (τ_membrane_) obtained from single exponential fits to the voltage response to −200pA current injection (C). *, #, and $ denote p<0.05 for differences between cell types, in GC across age groups and SGC across age groups, respectively by TW-ANOVA followed by post-hoc pairwise comparison (Supplementary Table 8).

