## Supplemental Tables for "Dendritic Morphology and Inhibitory Regulation Distinguish Dentate Semilunar Granule Cells from Granule Cells through Distinct Stages of Postnatal Development"

**Supplementary Table 1: Definitions of Somatic Parameters**

| SOMATIC PARAMETERS | | | |
|---|---|---|---|
| **Parameter** | **Acronym** (Suppl. Fig. 3) | **Equations** | **Definitions: MBF BIOSCIENCE** https://www.mbfbioscience.com/help/neurolucida_explorer/Content/Analyze/Branched%20Structure/neuronSumm.htm?Highlight=cell%20body%20details |
| *Soma Volume* | Svol | | Calculations for the cell body were accomplished by dividing the soma into stacks with equal thickness and quantifying the summed volume of the stacks. |
| *Surface Aea of soma* | Sarea | *[(perimeter Contour 1 + perimeter Contour 2 + ... + perimeter Contour n)/n] * thickness* | Addition of outer surface area of stack of all the contours. |
| *Soma Width* | Swith | | Measure of the largest distance in soma parallel to the hilar border |
| **Somatic parameters related to largest soma cross section based on highest perimeter** | | | |
| *Contour* | Cont | | Each slice division of the soma |
| *Feret Max* | Feret.X | | The largest dimensions of the contour as if a caliper was used to measure across the contour. |
| *Aspect ratio* | S_Asp_Rt | | This is a 2D parameter of soma contour defined as the ratio of its minimum diameter to its maximum diameter. [Range of values is 0-1. A circle has an aspect ratio of 1] |
| *Compactness* | S_Cmpct | $\sqrt{\frac{4}{\pi} * AREA}$/ Max Diameter | 2D parameter describes the relationship between the area and the maximum diameter. [Range of values is 0 to 1. A circle is the most compact shape with compactness = 1]. |
| *Form factor* | S_Form | $4\pi * AREA/$ (perimeter)$^2$ | The form factor differs from the compactness by considering the complexity of the perimeter of the object. As the contour shape approaches that of a perfect circle, this value approaches a maximum of 1.0. |
| *Roundness* | S_Rnd | [Compactness]$^2$ | Roundness is the square of the compactness to enable discrimination of objects with small compactness values. |
| *Convexity* | S_xity | Convex perimeter/Perimeter | A completely convex object does not have indentations, and has a convexity value of 1 (e.g., circles, ellipses, and squares). Concave objects have convexity values less than 1. |
| *Solidity* | S_solid | Area/Convex area | Solidity is the area of the contour divided by the convex area.<br>The area enclosed by a 'rubber band' stretched around a contour is called the convex area.<br>• Circles, squares, and ellipses have a solidity of 1.<br>• Indentations in the contour take area away from the convex area, decreasing the actual area within the contour.<br>Note that it is possible to have contours with low convexity and high solidity, and vice versa. |
| *Perimeter* | Speri | *Perimeter +/- (Perimeter Error Coefficient * Max distance from contour boundary in microns)* | Refers to the length of the contour for either open or closed contours.<br>The length takes the Z positions of the coordinates into account. |

**Supplementary Table 2: Definitions of Dendritic Parameters**

| **Parameter** | **Acronym** | **Equations** | **Definitions:** MBF BIOSCIENCE- https://www.mbfbioscience.com/help/neurolucida_explorer/Content/Analyze/Branched%20Structure/neuronSumm.htm?Highlight=cell%20body%20details |
|---|---|---|---|
| *Centrifugal Ordering and Branch orders* | | | The centrifugal ordering scheme is used to assign branch order to a tree. The segment that begins at the origin of the dendrite is assigned the branch order **1**. The branches that connect to that segment are assigned the branch order **2** and so on until all branches are assigned a value. |
| *Tortuosity* | Tort | Tortuosity = [Distance along process] / [Straight line distance] | The smallest tortuosity possible is 1—This is a straight path. Tortuosity increases as the segment assumes a more complex path to reach its destination. Tortuosity allows segments of different lengths to be compared in terms of the complexity of the paths they take. |
| *Primary* | prim | | Number of dendrites originating directly from soma. |
| *Primary Secondary, tertiary, quaternary Dendritic segments* | x1segm, x2segm, x3segm & x4segm | | Number of dendritic segments in first, second, third and fourth order dendrites respectively. Note X1SEGM is the same as PRIM |
| *Primary, Secondary tertiary, length* | x1lth, x2lth, x3lth & x4lth | | Summed lengths of all segments in a given dendritic order |
| *Totals terminals, Primary Secondary, tertiary, quaternary Terminals* | end, x1end, x2end, x3end & x4end | | Number of distal ends or terminals of dendritic segments that don't divide further |
| *Dendritic Length* | D_Lth | x1lth + x2lth + x3lth……. | Total length of all the dendrites |
| *Dendritic Mean Length* | D_MnLth | D_Lth / no. of primary dendrites | Length per dendritic tree arising from soma |
| *Angle* | ang | | Two dimensional measurement of angle subtended by the most distal of the most distant dendrites at the center of soma |
| *Space volume of cell* | D_Area | Convex Hull | Volume of the space occupied by the dendritic arbors of the cell |
| *Surface Area of space occupied by cell* | D_Volume | Convex Hull | Surface area of the space occupied by the dendritic arbors of the cell |
| *Dendritic Complexity* | D_Cmplx | [Sum of the terminal orders + Number of terminals] * [Total dendritic length / Number of primary dendrites] | A parameter that normalizes and compares distribution of dendrites among fundamentally different neurons |
| *Maximum Order* | MxOrdr | | Maximum order up to which dendritic segments reach. |

**Supplementary Table 3: Mean and SEM of Anatomical Parameters**

| Parameter Tested | Cell Type | Infant ( Mean ± SEM) | Adolescent ( Mean ± SEM) | Adult ( Mean ± SEM) |
|---|---|---|---|---|
| Soma Width (W) 2D XY plane | GC | 13.23 ± 1.42 | 11.7 ± 1.42 | 10.125 ± 1.74 |
| | SGC | 14.52 ± 1.01 | 14.28 ± 1.56 | 13.11 ± 1.32 |
| Soma Ratio W/H | GC | 0.78 ±0.11 | 0.76 ± 0.11 | 0.68 ± 0.14 |
| | SGC | 0.99 ± 0.08 | 1.07 ± 0.13 | 1.17 ± 0.11 |
| Number of primary dendrites | GC | 1.83 ± 0.41 | 1.33 ± 0.41 | 1.25 ± 0.5 |
| | SGC | 3.42 ± 0.29 | 3.5 ± 0.41 | 3 ± 0.38 |
| Dendritic – Angle 2D | GC | 59.06 ± 9.54 | 56.26 ± 9.54 | 56.25 ± 11.69 |
| | SGC | 100.75 ± 6.75 | 115.44 ± 9.54 | 90.78 ± 8.83 |
| First Order Nodes | GC | 1.5 ± 0.39 | 1.33 ± 0.39 | 1.25 ± 0.48 |
| | SGC | 2.83 ± 0.27 | 3.17 ± 0.39 | 2.57 ± 0.36 |
| Second Order Nodes | GC | 2.33 ± 0.53 | 2.5 ± 0.53 | 2.0 ± 0.65 |
| | SGC | 3.67 ± 0.37 | 4.33 ± 0.53 | 3.57 ± 0.49 |
| Second Order Segments | GC | 3.33 ± 0.77 | 3.0 ± 0.77 | 2.75 ± 0.94 |
| | SGC | 5.92 ± 0.54 | 6.67 ± 0.77 | 5.14 ± 0.71 |
| Third Order Segments | GC | 4.67 ± 0.96 | 5.0 ± 0.96 | 3.5 ± 1.8 |
| | SGC | 7.42 ± 0.68 | 8.17 ± 0.96 | 7.28 ± 0.89 |
| Dendrite -length | GC | 1797.94 ± 296.38 | 2543.38 ± 296.38 | 2252.62 ± 36 |
| | SGC | 1757.42 ± 209.57 | 2612.96 ± 296.38 | 2593.9 ± 274.4 |
| Dendritic complexity | GC | 85756.7 ± 29347.5 | 187443.47 ± 29347.49 | 209155.77 ± 35943.19 |
| | SGC | 36503.66 ± 20751.81 | 69192.72 ±29347.5 | 64871.99 ± 27170.5 |
| Convex Hull 3D Volume | GC | 481432.8 ± 373662.5 | 1492612.5 ± 373662.51 | 1442064.5 ± 457641.23 |
| | SGC | 909250 ± 264219.3 | 1778502.0 ± 409326.77 | 2902048.0 ± 345944.26 |
| Convex Hull Surface Area | GC | 53749.7 ± 14330.2 | 94378.55 ± 14330.19 | 98233.650 ± 17550.83 |
| | SGC | 82789.31 ± 10132.97 | 148579.2 ± 15697.93 | 159553.66 ± 13267.18 |
| Total Ends | GC | 15.17 ± 1.76 | 16.17 ± 1.76 | 14.0 ± 2.16 |
| | SGC | 15.5 ± 1.25 | 19.33 ± 1.76 | 14.57 ± 1.63 |
| Nodes Total | GC | 13.83 ± 2.05 | 17.000 ± 2.05 | 16.75 ± 2.51 |
| | SGC | 12.33 ± 1.45 | 16.67 ± 2.05 | 13.71 ± 1.9 |

| | | | | |
|---|---|---|---|---|
| Soma Surface Area | GC | 1044.31 ± 236.86 | 870.54 ± 236.86 | 657.14 ± 290.09 |
| | SGC | 1178.09 ± 167.49 | 1191.94 ± 259.47 | 784.25 ± 236.86 |
| Dendritic Tortuosity | GC | 1.08 ± 0.01 | 1.06 ± 0.01 | 1.1 ± 0.015 |
| | SGC | 1.06 ± 0.01 | 1.077 ± 0.01 | 1.09 ± 0.01 |
| Total Dendritic Terminals | GC | 15.17 ± 1.76 | 16.17 ± 1.76 | 14.0 ± 2.16 |
| | SGC | 15.5 ± 1.25 | 19.33 ± 1.76 | 14.57 ± 1.63 |

### Supplementary Table 4 : Statistical Summary of Anatomical Parameters

| Parameter Tested | Age Effect (F, p) | Cell Type Effect (F, p) | Tests |
|---|---|---|---|
| Soma Width (W) 2D XY plane | NS $F(2, 33) = 1.329$ $p = 0.278$ | **Sig** $F(1,33) = 4.457$ $P = 0.042$ | TW-ANOVA Bonferroni t-test Shapiro Wilk (Normality test) = Passed Equal Variance Test = Passed |
| Soma Ratio W/H | NS $F(2, 33) = 0.0665$ $p = 0.936$ | **Sig** $F(1,33)= 13.106$ $p < 0.001$ | TW-ANOVA Bonferroni t-test Shapiro Wilk (Normality test) = Passed Equal Variance Test = Passed |
| Number of primary dendrites | NS $F(2,34)= 0.771$ $p = 0.470$ | **Sig** SGC > GC $F(1,34)=30.695$ $p < 0.001$ | TW-ANOVA Bonferroni t-test Shapiro Wilk (Normality test) = Failed Equal Variance Test = Passed |
| Dendritic – Angle 2D | NS $F(2,34)= 0.768$ $p = 0.472$ | **Sig** SGC >GC $F(1,34)=34.371$ $p < 0.001$ | TW-ANOVA Bonferroni t-test Shapiro Wilk (Normality test) = Failed Equal Variance Test = Passed |
| First Order Nodes | NS $F(2,34)= 0.376$ $p = 0.689$ | **Sig** $F(1,34) = 22.712$ $p < 0.001$ | TW-ANOVA Bonferroni t-test Shapiro Wilk (Normality test) = Passed Equal Variance Test = Passed |
| Second Order Nodes | NS $F(2,34)= 0.700$ $p = 0.503$ | **Sig** $F(1,34) = 13.748$ $p < 0.001$ | TW-ANOVA Bonferroni t-test Shapiro Wilk (Normality test) = Passed Equal Variance Test = Passed |
| Second Order Segments | NS $F(2,34)= 0.668$ $p = 0.519$ | **Sig** SGC > GC $F(1,34)=21.741$ $p < 0.001$ | TW-ANOVA Bonferroni t-test Shapiro Wilk (Normality test) = Passed Equal Variance Test = Passed |
| Third Order Segments | NS $F(2,34)= 0.706$ $p = 0.501$ | **Sig** SGC > GC $F(1,34)=17.441$ $p < 0.001$ | TW-ANOVA Bonferroni t-test Shapiro Wilk (Normality test) = Passed Equal Variance Test = Passed |
| Dendrite -length | **Sig** $F(2,34)=4.817$ $p = 0.014$ | NS $F(1,34)=0.267$ $p = 0.609$ | TW-ANOVA Bonferroni t-test Shapiro Wilk (Normality test) = Passed Equal Variance Test = Passed |
| Dendritic complexity | **Sig** $F(2,34)= 4.597$ $p = 0.017$ | **Sig** $F(1,34) = 19.270$ $p < 0.001$ | TW-ANOVA Bonferroni t-test Shapiro Wilk (Normality test) = Failed Equal Variance Test = Failed |
| Convex Hull 3D Volume | **Sig** $F(2, 33) = 8.714$ $p < 0.001$ | **Sig** $F(1,33)= 5.587$ $p = 0.024$ | TW-ANOVA Bonferroni t-test Shapiro Wilk (Normality test) = Failed Equal Variance Test = Passed |
| Convex Hull Surface Area | **Sig** $F(2, 33) = 12.024$ $p < 0.001$ | **Sig** $F(1,33)= 16.801$ $p = <0.001$ | TW-ANOVA Bonferroni t-test Shapiro Wilk (Normality test) = Passed Equal Variance Test = Passed |
| Total Ends | NS $F(2,34)= 1.940$ $p = 0.159$ | NS $F(1,34)= 0.909$ $p = 0.347$ | TW-ANOVA Shapiro Wilk (Normality test) = Passed Equal Variance Test = Passed |
| Nodes Total | NS $F(2, 33) = 1.959$ $p = 0.156$ | NS $F(1,33)= 0.965$ $p = 0.333$ | TW-ANOVA, Shapiro Wilk (Normality test) = Passed Equal Variance Test = Passed |
| Soma Surface Area | NS $F(2, 32) = 1.409$ $p = 0.259$ | NS $F(1,32)= 0.975$ $p = 0.331$ | TW-ANOVA Bonferroni t-test Shapiro Wilk (Normality test) = Failed Equal Variance Test = Passed |
| Dendritic Tortuosity | NS $F(2,34)= 3.020$ $p = 0.062$ | NS $F(1,34)= 0.215$ $p = 0.646$ | TW-ANOVA Shapiro Wilk (Normality test) = Passed Equal Variance Test = Passed |
| Total Dendritic Terminals | NS $F(2,34)= 1.940$ $p = 0.159$ | NS $F(1,34) = 0.909$ $p =0.347$ | TW-ANOVA Shapiro Wilk (Normality test) = Passed Equal Variance Test = Passed |
| NS: Not Significant; Sig: Significant | | | |

**Supplementary Table 5: Developmental profile of tonic and synaptic inhibition in SGCs and GCs**

| Cell Property | Cell Type | Infant (P11-13) | Adolescent (P28-42) | Adult (P>120) | p-value |
|---|---|---|---|---|---|
| **sIPSC Frequency (Hz) FIGURE 4** | GC Mean±SEM | 10.6±1.3 (n=6/4) | 17.5±1.5 (n=13/6) | 15.3±2.2 (n=14/7) | Effect of cell type (p<0.05) Effect of age (p<0.05) Two-way ANOVA |
| | GC Median (IQR) | 6.4 (2.565 -13.7) | 11.7 (6.0- 23.1) | 8.5 (3.4 – 19.0) | |
| | SGC Mean±SEM | 16.5±2.0 (n=12/7) | 26.2±1.8 (n=11/9) | 15.7±1.2 (n=14/7) | |
| | SGC Median (IQR) | 12.8 (6.2-23.9) | 19.6 (10.6-37.8) | 11.3 (5.5-23.8) | |
| **sIPSC Amplitude (pA) FIGURE 5** | GC Mean±SEM | 48.5±8.3 (n=6/4) | 31.4±3.1 (n=13/6) | 27.0±1.3 (n=14/7) | Effect of cell type (p>0.05) Effect of age (p<0.05) Two-way ANOVA |
| | GC Median (IQR) | 38.5 (24.7-65.6) | 26.294 (44.3-18.8) | 25.5 (37.7-18.2) | |
| | SGC Mean±SEM | 51.4±6.1 (n=12/7) | 24.7±2.7 (n=11/9) | 21.1±2.0 (n=14/7) | |
| | SGC Median (IQR) | 42.1 (26.9 – 84.3) | 27.6 (19.0- 39.8) | 20.385 (15.9-30.2) | |
| **sIPSC Rise Time (ms) FIGURE 5** | GC Mean±SEM | 0.34±0.04 (n=6/4) | 0.20±0.02 (n=13/6) | 0.24±0.03 (n=14/7) | Effect of cell type (p=0.372) Effect of age (p=0.004) Two-way ANOVA |
| | GC Median (IQR) | 0.3 (0.25-0.3) | 0.2 (0.2-0.2) | 0.2 (0.2-0.25) | |
| | SGC Mean±SEM | 0.23±0.03 (n=12/7) | 0.24±0.03 (n=11/9) | 0.39±0.03 (n=14/7) | |
| | SGC Median (IQR) | 0.2 (0.2-0.25) | 0.25 (0.2-0.25) | 0.4 (0.25-0.5) | |
| **sIPSC $\tau_{decay\text{-}WT}$ (ms) FIGURE 5** | GC Mean±SEM | 6.00±0.82 (n=6/4) | 3.69±0.60 (n=13/6) | 2.88±0.58 (n=14/7) | Effect of cell type (p=0.379) Effect of age (p=0.635) Two-way ANOVA |
| | GC Median (IQR) | 5.94 (4.47-7.61) | 3.95 (2.98-4.37) | 2.75 (2.59-3.2) | |
| | SGC Mean±SEM | 3.38±0.72 (n=12/7) | 4.46±0.65 (n=11/9) | 6.16±0.54 (n=14/7) | |
| | SGC Median (IQR) | 3.65 (3.37-5.83) | 3.16 (2.93-3.76) | 6.52 (3.22-10.12) | |
| **sIPSC Area (pA.ms) FIGURE 4** | GC Mean±SEM | 421.24±42.45 (n=6/4) | 149.68±31.15 (n=13/6) | 114.85±30.02 (n=14/7) | Effect of cell type (p=0.658) Effect of age (p<0.001) Two-way ANOVA |
| | GC Median (IQR) | 399.13 (343.62-473.59) | 134.29 (125.68-170.72) | 109.43 (94.98-135.78) | |
| | SGC Mean±SEM | 376.33±37.44 (n=12/7) | 164.70±31.15 (n=11/9) | 182.03±31.15 (n=14/7) | |
| | SGC Median (IQR) | 312.06 (235.37-450.9) | 121.44 (100.51-135.01) | 149.56 (119.81-217.79) | |
| **Charge Transfer per second (Area X Frequency)** | GC Mean±SEM | 4928.22±569.42 (n=6/4) | 2560.07±244.13 (n=13/6) | 1677.19±200.58 (n=14/7) | Effect of cell type (p=0.88) Effect of age (p<0.001) |
| | GC Median (IQR) | 4572.97 (4103.77-4747.09) | 2296.18 (1855.34-3360.22) | 1498.03 (1019.28-2391.51) | |
| | SGC Mean±SEM | 6568.51±1001.62 (n=12/7) | 2953.89±488.951 (n=11/9) | 2803.12±566.71 (n=14/7) | |

| | | | | | |
|---|---|---|---|---|---|
| **SUP. FIGURE 5** | SGC Median (IQR) | 5440.3 (4894.26-8106.04) | 2726.73 (2030.29-3500.48) | 2383.69 (1814.21-2793.8) | Two-way ANOVA on Median values |
| **Extrasynaptic Tonic GABA Current Amplitude (pA) FIGURE 6** | GC Mean±SEM | 3.96±1.7 (n=5/4) | 10.2±1.6 (n=13/6) | 5.4±1.4 (n=10/7) | Effect of cell type (p=0.147) Effect of age (p<0.001) Two-way ANOVA |
| | SGC Mean±SEM | 2.4±0.8 (n=9/5) | 17.1±1.7 (n=9/7) | 4.1±1.3 (n=10/6) | |
| **Input Resistance (MΩ) SUP. FIGURE 7** | GC Mean±SEM | 338±41.4 (n=6/4) | 245±21.5 (n=7/5) | 160.3±31.5 (n=5/3) | Effect of cell type (p=0.355) Effect of age (p=0.878) Two-way ANOVA |
| | GC Median (IQR) | 315.6 (164.8-348.4) | 268.4 (107.6-345.6) | 150.2 (135.1-205.7) | |
| | SGC Mean±SEM | 164.5±17.4 (n=7/4) | 168.3±14.5 (n=11/7) | 208.9±38.7 (n=7/4) | |
| | SGC Median (IQR) | 168 (135.5-191.8) | 178.7 (119.8-204.7) | 197.5 (111.5-309.1) | |
| **Membrane Tau SUP. FIGURE 7** | GC Mean±SEM | 14.5±2.1 (n=6/4) | 12.9±1.4 (n=7/5) | 8.1±0.7 (n=5/3) | Effect of cell type (p=0.449) Effect of age (p=0.686) Two-way ANOVA |
| | GC Median (IQR) | 14.2 (8.9-19.3) | 11.5 (9.5-16.6) | 7.9 (6.7-9.5) | |
| | SGC Mean±SEM | 9.3±0.4 (n=7/4) | 10.9±1.1 (n=11/7) | 20.3±5.8 (n=7/4) | |
| | SGC Median (IQR) | 9.9 (7.7-10.1) | 10.4 (7.6-12.6) | 11.7 (10.9-34.3) | |
| Note: n values were represented as (n=number of cells/number of rats); GCL- Granule Cell layer; ANOVA – Analysis of Variance | | | | | |

**Supplementary Table 6: Pairwise statistical analysis of physiological parameters**

| | A vs B (Former A and Latter B) | sIPSC Frequency Kruskal-Wallis Dunn’s Method (DOR) | sIPSC Amplitude Kruskal-Wallis Dunn’s Method (DOR) | TONIC TW-ANOVA S-W – Failed Benferroni Pairwise t-test (p) |
|---|---|---|---|---|
| **SGC** | Infant vs Adolescent | **Sig** B>A (258.378) | **Sig** A>B (339.820) | **Sig** B>A (<0.001) |
| | Infant vs Adult | NS (34.449) | **Sig** A>B (559.275) | NS (1.0) |
| | Adolescent vs Adult | **Sig** A>B (292.826) | **Sig** A>B (219.455) | **Sig** A>B (<0.001) |
| **GC** | Infant vs Adolescent | **Sig** B>A (310.180) | **Sig** A>B (269.107) | **Sig** B>A (0.035) |
| | Infant vs Adult | **Sig** B>A(150.122) | **Sig** A>B(319.609) | NS (1.0) |
| | Adolescent vs Adult | **Sig** A>B (160.057) | NS (50.502) | **Sig** A>B (0.043) |
| **GC vs SGC** | Infant | **Sig** B>A (328.787) | NS (65.173) | NS (0.53) |
| | Adolescent | **Sig** B>A (276.985) | NS (5.540) | **Sig** B>A (<0.001) |
| | Adult | **Sig** B>A (144.216) | **Sig** A>B (174.493) | NS (0.52) |
| | TW-ANOVA (Two Way Analysis of Variance), S-W (Shapiro-Wilk Normality test), DOR – Difference of Ranks), sIPSCs Failed Normality (S-W Test) for both amplitude and frequency | | | |

**Table 7: Pairwise statistical analysis of sIPSC kinetics parameters**

| | A vs B (Former A and Latter B) | sIPSC Rise Time (ms) TW-ANOVA S-W – Failed Benferroni Pairwise t-test (p) | sIPSC Decay Tau wt. TW-ANOVA S-W – Failed Benferroni Pairwise t-test (p) | sIPSC Charge transfer (pA.sec) TW-ANOVA S-W – Failed Benferroni Pairwise t-test (p) | sIPSC cumulative Charge Transfer in one Sec (pA.sec) TW-ANOVA S-W – Failed Benferroni Pairwise t-test (p) |
|---|---|---|---|---|---|
| **SGC** | Infant vs Adolescent | NS (p=1.00) | NS (p=0.15) | **Sig(A>B)** (p<0.001) | **Sig(A>B)** (p<0.001) |
| | Infant vs Adult | **Sig(B>A)** (p<0.001) | **Sig(B>A)** (p=0.009) | **Sig(A>B)** (p<0.001) | **Sig(A>B)** (p<0.001) |
| | Adolescent vs Adult | **Sig(B>A)** (p=0.001) | NS (p=0.81) | NS (p=1.00) | NS (p=1.0) |
| **GC** | Infant vs Adolescent | **Sig(A>B)** (p=0.01) | NS (p=0.08) | **Sig(A>B)** (p<0.001) | **Sig(A>B)** (p<0.001) |
| | Infant vs Adult | NS (p=0.164) | **Sig(B>A)** (p=0.008) | **Sig(A>B)** (p<0.001) | **Sig(A>B)** (p<0.001) |
| | Adolescent vs Adult | NS (p=1.00) | NS (p=1.0) | NS (p=1.0) | NS (p=1.0) |
| **GC vs SGC** | Infant | **Sig(A>B)** (p=0.25) | **Sig(A>B)** (p=0.02) | NS (p=0.43) | NS (p=0.06) |
| | Adolescent | NS (p=0.43) | NS (p=0.387) | NS (p=0.73) | NS (p=0.06) |
| | Adult | **Sig(B>A)** (p=0.001) | **Sig(B>A)** (p<0.001) | NS (p=0.12) | NS (p=0.06) |
| TW-ANOVA (Two Way Analysis of Variance), S-W (Shapiro-Wilk Normality test) failed for all the above parameters | | | | | |

**Table 8: Pairwise statistical analysis of sIPSC intrinsic physiology.**

| | **A vs B (Former A and Latter B)** | **Input Resistance TW-ANOVA S-W – Passed Benferroni Pairwise t-test (p)** | **Membrane Tau TW-ANOVA S-W – Failed Benferroni Pairwise t-test (p)** |
|---|---|---|---|
| **SGC** | Infant vs Adolescent | NS (p=1.0) | NS (p=1.0) |
| | Infant vs Adult | **Sig(B>A)** (p=0.03) | **Sig(B>A)** (p=0.02) |
| | Adolescent vs Adult | **Sig(B>A)** (p=0.01) | **Sig(B>A)** (p=0.04) |
| **GC** | Infant vs Adolescent | NS (p=1.0) | NS (p=1.0) |
| | Infant vs Adult | **Sig(A>B)** (p=0.005) | NS (p=0.46) |
| | Adolescent vs Adult | **Sig(A>B)** (p=0.018) | NS (p=0.82) |
| **GC vs SGC** | Infant | **Sig(A>B)** (p=0.006) | NS (p=0.17) |
| | Adolescent | **Sig(A>B)** (p=0.009) | NS (p=0.58) |
| | Adult | **Sig(B>A)** (p=0.003) | **Sig(B>A)** (p=0.01) |
| | TW-ANOVA (Two Way Analysis of Variance), S-W (Shapiro-Wilk Normality test) | | |
