## Supplementary Figures for "Dendritic Morphology and Inhibitory Regulation Distinguish Dentate Semilunar Granule Cells from Granule Cells through Distinct Stages of Postnatal Development"

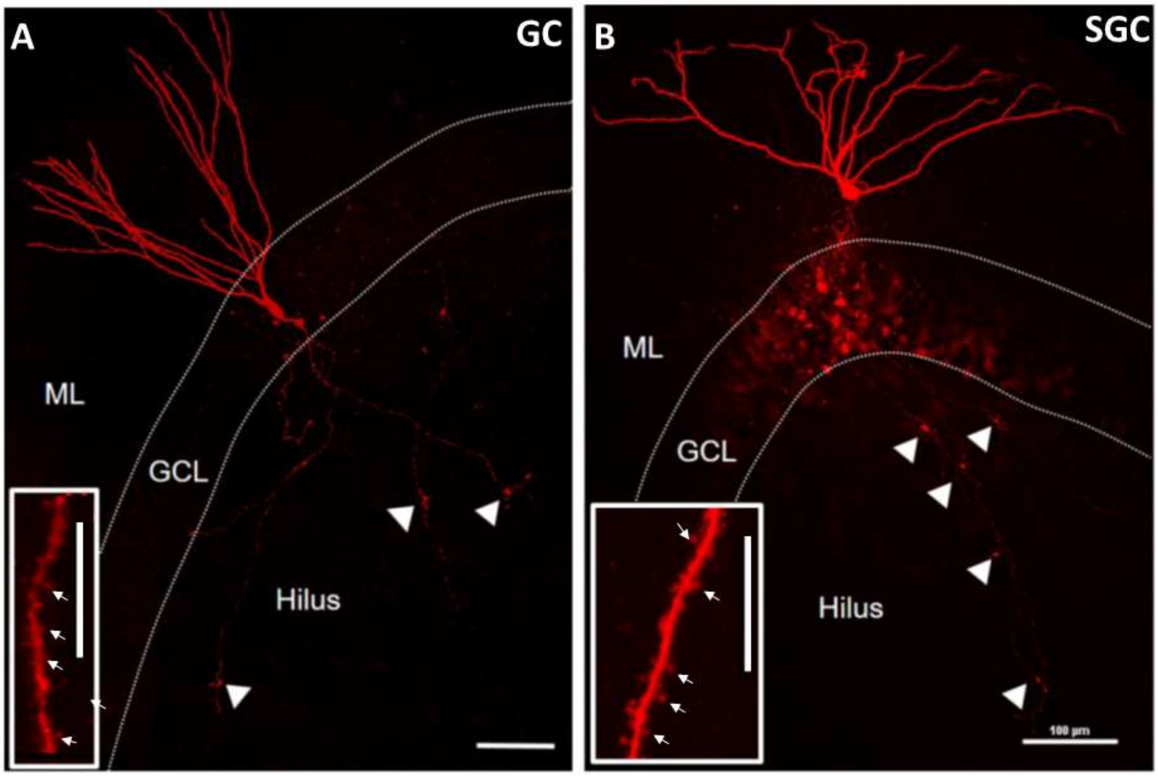


**Supplementary Figure 1: Representative images of a GC and SGC.** Images of a typical GC (A) and SGC (B) illustrate the somatic location, dendritic arbor, with high density of spines (insets, white arrows) and axons with boutons (white arrow heads) targeting CA3 used by experimenter to classify SGCs and GCs. Scale bar: 100µm; Inset scale bar: 20 µm.

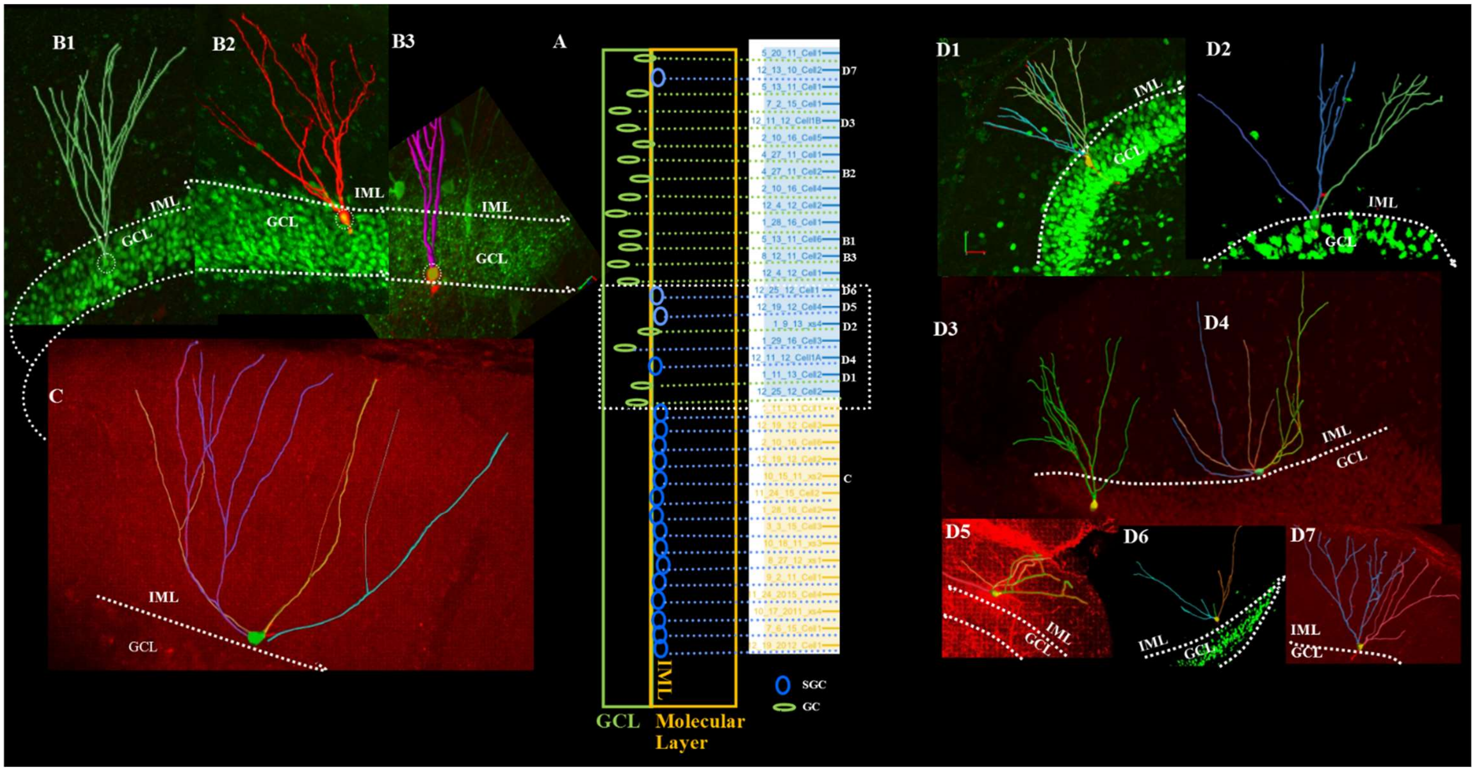


**Supplementary Figure 2: Distribution of somatic location of cells included in morphometric analysis.** A. Schematic of the somatic location in the GCL (green box) and IML (orange box) overlaid on the cluster analysis distribution illustrates SGCs in blue and GCs in green. $B_{1-3}$: Montage of biocytin filled images of representative GCs (labeled as $B_{1-3}$ in A) with GCL demarcated by dotted lined identified based on Prox-1 ($B_{1-2}$) or parvalbumin ($B_3$) immunostaining. C. Representative SGC reconstruction in IML overlaid on image of the slice. $D_{1-7}$. Example of biocytin filled and reconstructed GCs and SGCs labeled as $D_{1-7}$ in panel A. Note that the SGCs illustrated here clustered with putative GCs and are included in the white dotted area in A. All images were obtained a 20X magnification and reconstructed. Note the difference in soma shape of cells classified as GCs ($D_{1-3}$) versus those classified as SGCs($D_{4-7}$) by investigator. $D_7$. Note the increased dendritic complexity compared to putative SGCs in $D_{4-6}$. GCL: Granule Cell Layer, IML: Inner Molecular Layer.

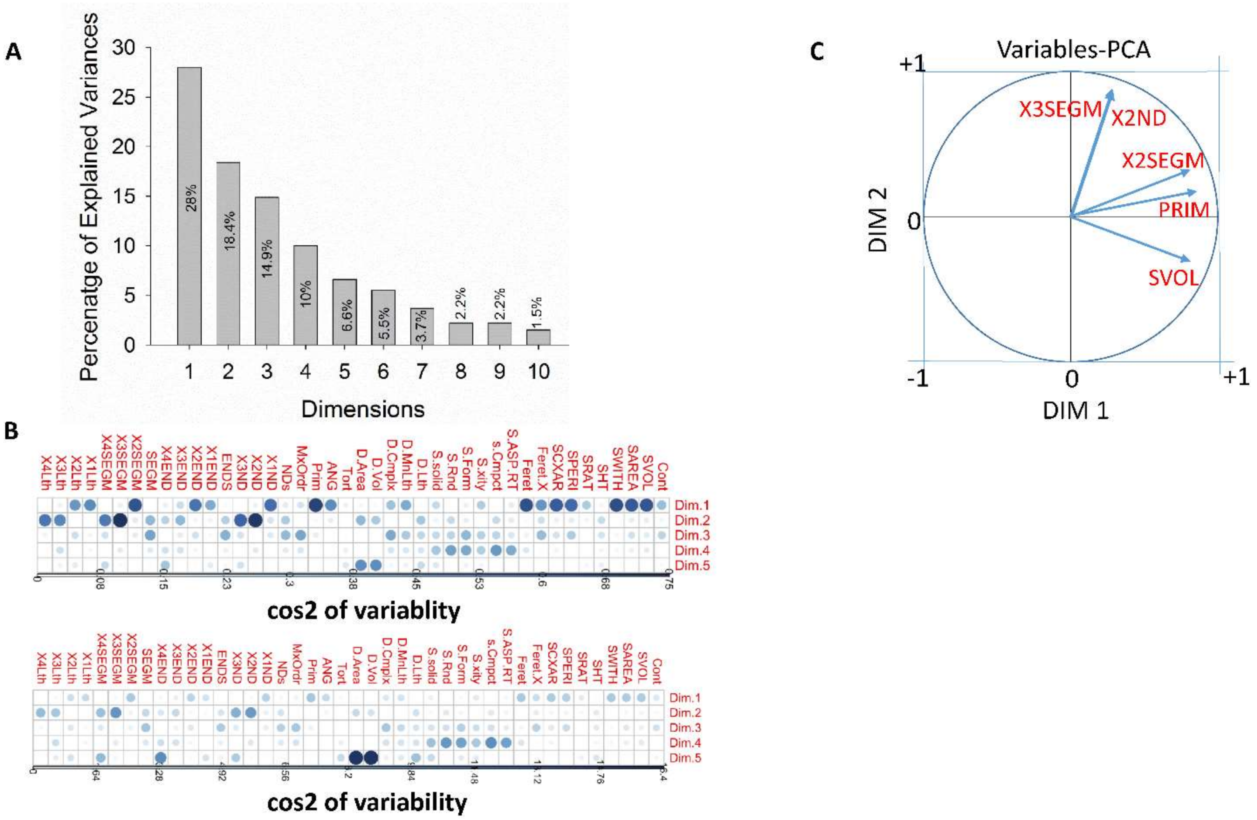


**Supplementary Figure 3: Analysis of Morphological Variables underlying Principal Components.** A. Histogram illustrates percentage of information retained by each dimension (principal component, PC). B. Factor maps illustrate the quality of representation of the morphometric variables measured by cos2 (square cosine, squared coordinates). The darker color indicates stronger contribution to variability to PC . C. Representation of the top five variables in the first two dimensions.

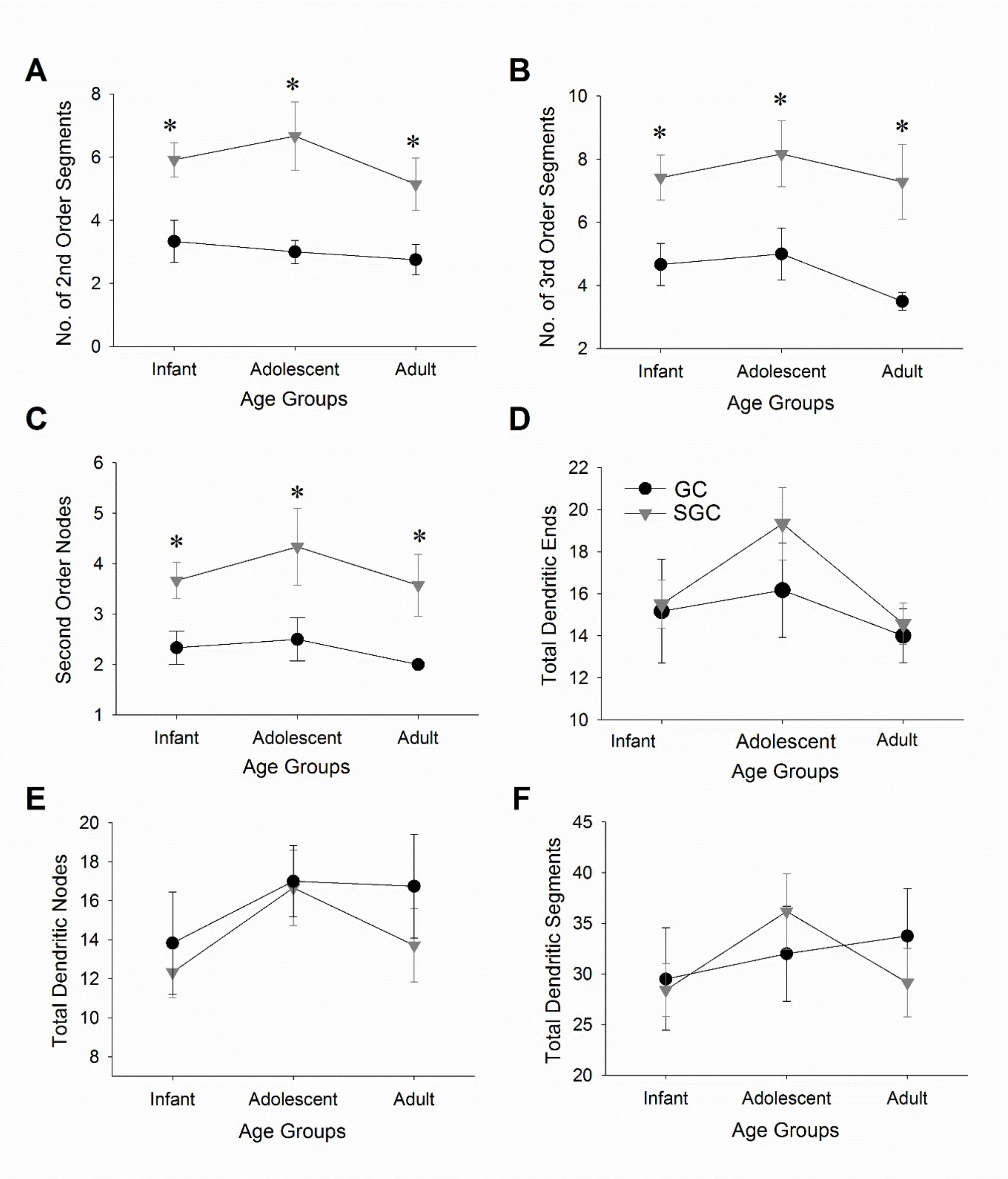


**Supplementary Figure 4: Comparison of morphometric parameters between GCs and SGCs at distinct developmental stages.** Summary plot of number of second (A) and third (B) order dendritic segments, second order nodes (C), total number of dendritic ends (D), total dendritic nodes (E) and total dendritic segments (F) in GCs and SGCs at the three age groups examined. * denotes $p<0.05$ for differences between cell types by TW-ANOVA followed by post-hoc pairwise comparison (Supplemental Table 3 &4). N= 6 infant, 6 adolescent and 4 adult GCs and 9 infant, 5 adolescent and 6 adult SGCs.

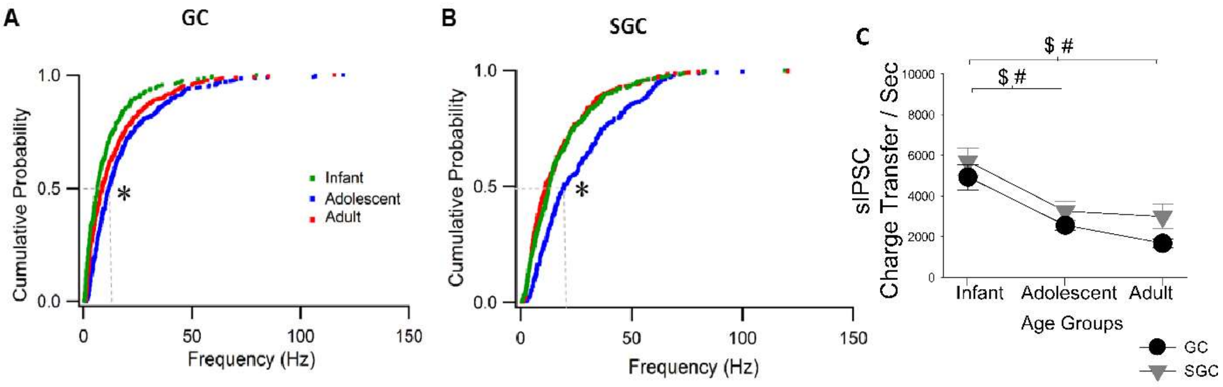


**Supplementary Figure 5: Cumulative plots of sIPSC frequency in GCs and SGCs through development.** Cumulative plots of sIPSC frequency comparing three age groups * denotes $p<0.05$ for differences between cell types by Kruskal-Wallis Test (Supplementary Table 6) in GCs (A) and SGCs (B). Summary plot of cumulative charge transfer over one second (C) at three developmental stages in both cell types. #, and $ denote $p<0.05$ for differences in GC across age groups and SGC across age groups, respectively by TW-ANOVA followed by post-hoc pairwise comparison.

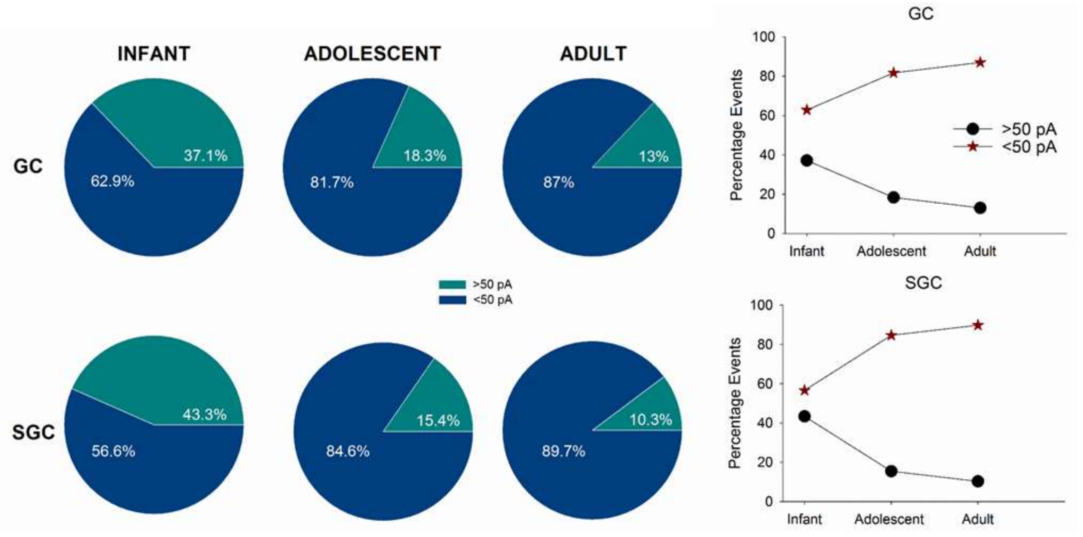


**Supplementary Figure 6: Developmental differences in distribution of sIPSC amplitudes in GCS and SGCs.** Pie chart distributions illustrating subjective percentage distribution of high peak amplitude (>50pA) and smaller peak amplitude (<50 pA) amplitude sIPSCs in GCs (top) and SGCs (bottom) across age groups.

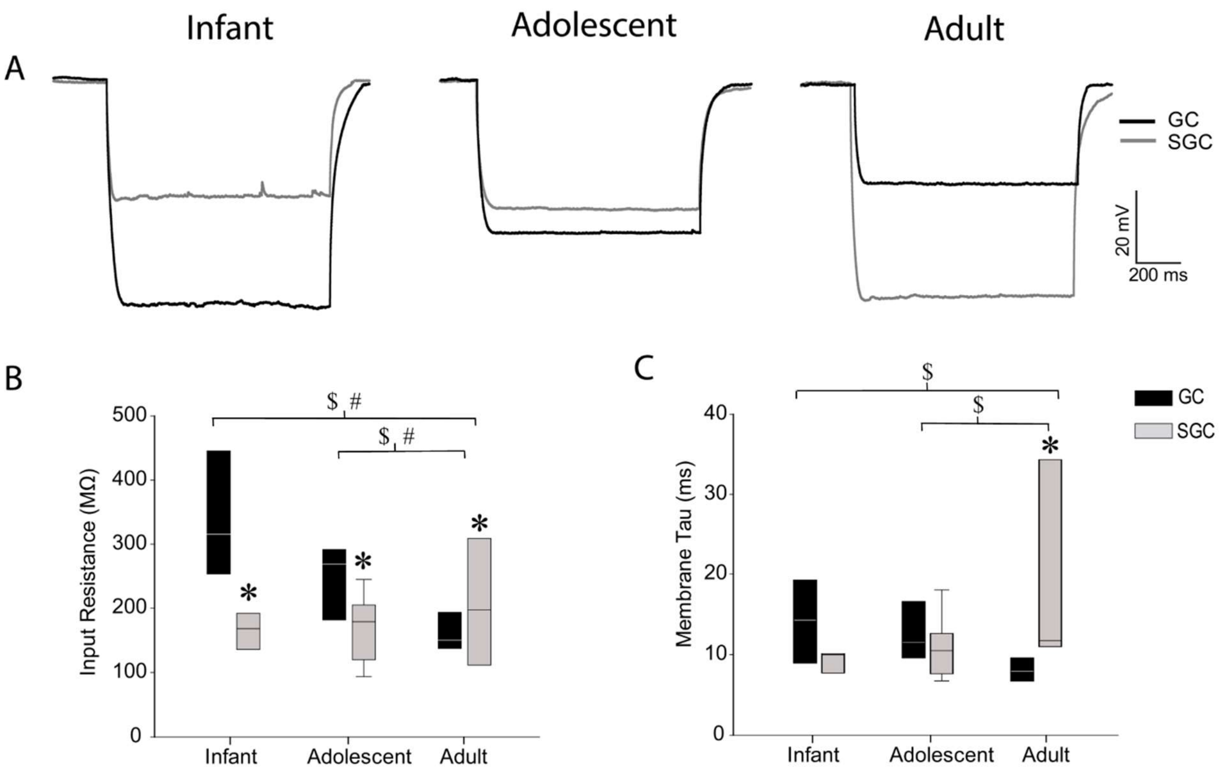


**Supplementary Figure 7: Analysis of developmental changes in passive membrane properties of GCs and SGCs.** Representative voltage traces in response to a -200pA current injection for one second in GCs and SGCs at the developmental stages under investigation reveal differences in $R_{in}$ (A). Summary of $R_{in}$ in the cell types (B). Summary plot of membrane time constant ($\tau_{membrane}$) obtained from single exponential fits to the voltage response to -200pA current injection (C). *, #, and $ denote $p<0.05$ for differences between cell types, in GC across age groups and SGC across age groups, respectively by TW-ANOVA followed by post-hoc pairwise comparison (Supplementary Table 8).
